# *In situ* structure of a dimeric hibernating ribosome from a eukaryotic intracellular pathogen

**DOI:** 10.1101/2022.04.29.490036

**Authors:** Mathew McLaren, Patricia Gil-Diez, Michail N. Isupov, Rebecca Conners, Lavinia Gambelli, Vicki Gold, Andreas Walter, Sean R. Connell, Bryony Williams, Bertram Daum

## Abstract

Translational control is an essential process for the cell to adapt to varying physiological or environmental conditions. To survive adverse conditions such as low nutrient levels, translation can be shut down almost entirely by inhibiting ribosomal function. Here we investigated eukaryotic hibernating ribosomes from the microsporidian parasite *Spraguea lophii in situ* by a combination of cryo-electron tomography (cryoET) and single particle cryoEM. We show that microsporidian spores contain ribosomes primed for host cell invasion and thus shed new light on the infection mechanism of this important pathogen. Prior to host infection, virtually all ribosomes are locked in the 100 S dimeric state, which appears to be formed by a unique dimerization mechanism that is distinct from its bacterial counterparts. Within the dimer, the hibernation factor MDF1 is bound within the E site, locking the L1 stalk in a closed conformation, and thus preventing the translation of mRNAs to polypeptides.

## Introduction

The ribosome is the central protein production hub of the cell and conserved throughout evolution. The process of protein translation is very energy expensive and can consume up to 40 % of all cellular energy^1^. Under adverse environmental or physiological conditions, a large proportion of the cell’s ATP consumption can be saved by down-regulating translation at the level of the ribosome^2^. This is achieved by various ribosomal inhibitors, a subset of which promote ribosomal hibernation^3^. These so-called hibernation factors inactivate ribosomes by two mechanisms; either by locking individual ribosomes in a state that is incompatible with translation or through promoting the formation of dimeric, 100 S ribosomes. Either mechanism prevents the dissociation of the small and large subunit, which is an essential prerequisite for mRNA loading and translation^1^. Moreover, ribosome dimerisation protects the ribosomes from degradation by RNases^4^.

The mechanisms of ribosomal hibernation are particularly well investigated in bacteria^4^. In stationary growth phases or under conditions of nutrient-starvation, two forms of hibernating ribosomes appear to co-exist. In *E. coli* these are a 70 S ribosome, inactivated by the hibernation factor RaiA, and a 100 S ribosome dimer, stabilized by the binding of up to two hibernation factors called ribosome modulation factor (RMF) and hibernation promoting factor (HPF)^2, 5, 6^. In combination, these factors block ribosome function by binding and occluding the decoding centre, the mRNA binding channel and the acceptor (A) and peptidyl (P) tRNA sites^6–8^. A recent cryoEM study of hibernating ribosomes from *E. coli* revealed a 100 S particle that, in addition to HPF and RMF, was also bound to the factor bS1, as well as a deacetylated tRNA in the E site. Here, bS1 is stabilized by RMF and together both factors sequester the anti-Shine-Dalgarno sequence of the 16 S rRNA. In addition, the E-tRNA is stabilised by HPF, which itself occludes the binding site for the mRNA as well as A- and P-site tRNAs^9^. Many other bacteria, such as *Thermus thermophilus*, do not express RMF and instead rely on an extended version of HPF, HPFlong, for 100 S ribosome formation^4^.

In eukaryotes, ribosomal hibernation has so far been structurally characterized at the single (80 S) ribosome level. In hibernating yeast ribosomes, ribosomal function is blocked by the protein Lso2, which sequesters the mRNA binding pocket as well as the polypeptide exit tunnel concomitantly^10^. In mammals, various types of hibernating ribosomes can be found. In hibernating ribosomes isolated from human cell culture, two silencing states appear to co-exist: a non-rotated/non-ratcheted state bound to the Lso2 homolog CCDC124, as well as EBP1 at the polypeptide exit site and a second rotated state, where CCDC124 is exchanged by SERBP1 and eEF2^10^. Whereas CCDC124 / SERBP1 and eEF2 are analogous in occupying the mRNA entry channel and blocking the A and P sites, EBP1 prevents the stalled ribosome’s futile interaction with proteins interacting with the nascent polypeptide chain, including the ribosome associated complex (RAC), signal recognition particle (SRP), secretory protein 61 (Sec61), and N-α -Acetyltransferase A (NatA)^10^.

An alternative mammalian hibernation mechanism has been discovered in ribosomes isolated from reticulocytes. Here, 80 S ribosomes are inactivated by binding interferon-related developmental regulator 2 (IFRD2). IFRD2 occupies the P and E sites and sequesters the mRNA with its alpha-helical C-terminus. At the same time, a deacylated tRNA is bound to a non-canonical site beyond the E site, called the Z site. It was hypothesized that stably bound deacetylated Z-tRNA may present a hallmark of stalled ribosomes under scenarios of amino acid depletion, or where translational factors are limiting^11^. Biochemical evidence and negative stain electron microscopy suggest that ribosomal dimerisation also occurs in rat cells during amino acid starvation^12^. However so far, no structure of a eukaryotic ribosome dimer is available.

By combining cryo electron tomography (cryoET) with single particle cryoEM, we investigated the structure of eukaryotic hibernating ribosome dimers in the microsporidian species *Spraguea lophii.* Microsporidia are single-celled eukaryotic intracellular parasites with species infecting almost all animal lineages^13^. They have the potential to cause debilitating disease in immunocompromised humans and severe deleterious impacts on food production^14^. Microsporidia begin their life cycle as dormant spores that need to enter and exploit the energy metabolism of a host cell in order to proliferate^13^. They achieve entry into host cells via a preformed, tightly coiled ‘polar tube’ (PT) that is rapidly expelled from the dormant spore on germination^15^. The PT then penetrates the host cell membrane and the spore content (sporoplasm) is rapidly transported down the tube into the host cell^15^. As the PT is usually not more than 150 nm wide, this presents a perfect opportunity to image hibernating ribosomes by cryoET *in situ*.

Single particle cryoEM of ribosomes isolated from the microsporidian species *Varimorpha necatrix*^16^*, Paranosema locustae*^17^ and *Encephalitozoon* cuniculi^18^ have revealed structures of monomeric ribosomes in a hibernating state. These studies confirm the highly reduced nature of microsporidian ribosomes that show a drastic loss of the expansion segments characteristic of eukaryotic rRNA, but that have largely retained the core set of typical ribosomal proteins. The complexes are similar in size to their bacterial counterparts and are thus designated as 70 S ribosomes^16, 17, 19^. Similar to yeast, the ribosomes of *P. locustae* were shown to be inhibited by the protein Lso2 which spans the mRNA decoding site and the large subunit tRNA binding site. In contrast, hibernating ribosomes of *V. necatrix* were bound to two inhibiting factors, called MDF1 (a conserved eukaryotic protein) and MDF2 (a protein only identified in *V. necatrix, N. ceranae* and *N. apis*). While MDF1 binds the E site of the SSU and stabilizes the ribosome in a conformation incompatible with translation, MDF2 blocks the P site as well as the polypeptide exit tunnel of the ribosome. In *E. cuniculi*, the ribosome was bound to MDF1 in the E-site; acting as a mimic of a deacylated tRNA molecule.

By investigating PTs ejected from the species *S. lophii* by cryoET and single particle analysis, we find that sporoplasm traversing the PT is densely packed with hibernating 100 S ribosome dimers, which so far have only been characterized in detail in bacteria. The conformation of the 100 S ribosomes is markedly distinct from any of the bacterial structures known so far. Through the study of this unique group of organisms, we present the first cryoEM evidence of ribosomal dimerisation as a mechanism for ribosomal hibernation in eukaryotes.

## Materials and Methods

### Spore preparation

To isolate spores of *S. lophii*, clusters of cysts were collected from monkfish (*Lophius piscatorius)* caught in the North Atlantic and landed in Devon (United Kingdom). Xenomas filled with microsporidia spores were removed from fish tissue manually. Samples were homogenized manually using a scalpel in PBS buffer until a suspension was obtained. The suspension was filtered through a 100 μm mesh sterile cell strainer (Fisher Scientific). The spores were cleaned by centrifugation in a 25-50-75-100 % Percoll gradient (Sigma) at 4°C and 3,240 x g for 1 h. Spores were collected and washed three times in sterile PBS. The sample was stored at 4°C with the addition of 10 μg/ml of ampicillin. The concentration of spores in the solution was determined using a haemocytometer.

### On-grid spore germination

*S. lophii s*pores in suspension were mixed in 1:1 proportion with 10 nm protein A-gold (Aurion, Wageningen, The Netherlands) as fiducials. 3 μl of the mixture was deposited on glow discharged R2/2 Copper 300 mesh holey carbon-coated support grids (Quantifoil, Jena, Germany) along with 1 μl of HEPES pH 10 and 1 μl of hydrogen peroxide 0.9 %. The grids were initially screened and optimised in negative stain. For cryoET, grids were plunge-frozen in liquid ethane using a Vitrobot Mark-IV (Thermo-Fisher, Eindhoven, NL) using variable blot times of 4-6 sec and a blot force of -1.

The grids were screened using a 120 kV Tecnai Spirit (Thermo Fisher Scientific). Tilt series were collected with a 300 kV Titan Krios microscope (Thermo Fisher Scientific) at the Electron Bio-imaging Centre (eBIC, Diamond Light Source, Harwell, UK), equipped with a K2 Summit direct electron detector (Gatan, Pleasanton, USA). The tilts were collected using the SerialEM software^20^ with a pixel size of 4.377 Å. A nominal defocus range of -4 μm to -6 μm was used. A tilt range of -60° to +60° with 2° steps in a dose-symmetric tilt scheme^21^ was used for collection with a total target dose of 120 e^-^/Å^2^. A total of 35 tomograms were collected, with six containing ribosomes that were used for sub-tomogram averaging. Data collection details are shown in Table 1.

**Table 1.** Cryo tomography data collection

|  |  |
| --- | --- |
| Acceleration voltage (kV) | 300 |
| Pixel size (Å) | 4.377 |
| Defocus range (µm) | -4 to -6 (0.5 step) |
| Frames per tilt | 10 |
| Dose per tilt (e<sup>-</sup>/Å<sup>2</sup>) | 1.90 |
| Tilt range (°) | -60 to +60 |
| Tilt step (°) | 2 |
| Total dose (e<sup>-</sup>/Å<sup>2</sup>) | 116 |

### Tilt series reconstruction and sub-tomogram averaging

Tilt series motion correction, alignment and reconstruction were performed using the IMOD package^22^. Particles were initially picked manually using IMOD, then CTF estimation and extraction were performed using the Relion STA pipeline^23^. A total of 5001 particles were selected. 2D and 3D classification steps and a refinement of the entire dimer were performed in Relion 3.1^24^ to observe the dimer structure and determine the heterogeneity of the ribosomes. A refined average was created using 2871 particles. The resolution was found at 20.8 Å using Gold Standard Fourier Shell Correlation (FSC).

For higher resolution averaging, 3D CTF correction and template matching were performed on six tomograms in emClarity^25^, using a map of the *S. lophii* ribosome lowpass filtered to 40 Å as a reference. After removal of false positives, 4590 particles were selected and used for sub-tomogram averaging. Iterative alignments and tilt series refinements were performed at 2x and 1x binning to achieve a Gold Standard FSC resolution of 10.3 Å of half of a dimer pair. The final map was reconstructed from the tilt series images with a maximum exposure of 100 electrons. To reconstruct the full dimer, the alignment was shifted by 200 Å and the mask/search region expanded to encompass both ribosomes. As with the single ribosome, iterative alignments and tilt series refinements were performed at 2x and 1x binning, with classification performed at 2x binning. For the full dimer map a Gold Standard FSC resolution of 15.2 Å was reached with 1284 particles.

### Ribosome purification

*S. lophii* spores were washed and resuspended in sterile BS100 buffer (25 mM HEPES, 100 mM potassium acetate, 15 mM magnesium acetate and 1 mM DTT). Spores were disrupted using a Fastprep-24 5G bead beater (MP Biomedicals, Fisher Scientific, Sweden) and 0.1 mm glass beads. Cell debris and non-disrupted spores were pelleted by centrifugation at 9,000 x g for 15 min at 4°C. The supernatant was collected and layered on top of a 30 % sucrose cushion in BS100 buffer and centrifuged at 206,000 x g for 3 h at 4°C. The supernatant was discarded, and the pellet was resuspended by gentle shaking for 1h at 4°C in BS100 buffer. The crude extract was layered over a continuous sucrose gradient (10 – 40 %) in BS100 buffer. The sample was cleared by centrifugation at 33,000 x g for 80 min at 4°C. Fractions were collected and analysed for RNA using a Nanodrop (Thermo Scientific) at 260 nm. Vivaspin columns (Sartorius, Germany) were used to remove the excess of sucrose and concentrate the sample to a concentration between 5-15 mg/ml.

### Single particle cryoEM

3 μl of purified ribosome sample was applied to Quantifoil 300 mesh R2/2 grids with an ultrathin 2 nm carbon layer and frozen in liquid ethane using a Vitrobot Mark IV (Thermo Fisher Scientific) at 15 °C, 100 % relative humidity, blot force -1 and blot time 4 s.

Three datasets were collected on a 200 kV Talos Arctica microscope (Thermo Fisher Scientific) with a K2 Summit direct electron detector at the Wolfson Bioimaging Centre, University of Bristol, UK. Datasets were collected using EPU (Thermo Fisher Scientific) with pixel sizes of 1.054 Å, two in counting mode and one in super-resolution (0.525 Å) with a defocus range of -0.8 μm to -2 μm. A final dataset was collected on a 300 kV Titan Krios microscope with a K3 direct electron detector at eBIC. The data were collected using EPU with a pixel size of 1.06 Å (0.53 Å super-resolution).

Motion correction, CTF estimation and particle picking were performed using Warp^26^. A total of 977,879 particles were picked from 26997 micrographs. A full breakdown of the datasets is shown in Table 2. All four datasets were initially processed separately and combined for the later refinement steps. Several iterations of 2D and 3D classification and refinement were performed in Relion 3.1^24^. The two maps used for model building were a three-body refinement of the large subunit, small subunit body and small subunit head (285,940 particles) and a refinement of the best class of a 3D classification with no alignment. The resolutions obtained for these maps were 2.26 Å (2.26 Å, 2.48 Å and 2.86 Å for LSU, SSU body and SSU head respectively) and 2.78 Å for the whole-ribosome map. The L1 stalk resolved using a tight mask around the region, with a 3D classification with no alignment. The best class was then refined to a global resolution of 3.07 Å. The local resolution range for this region was 3 - 6 Å, with the density of L1 stalk and tRNA having a resolution of 5-6 Å. Post-processing of the maps was performed using DeepEMhancer^27^.

**Table 2.** Single particle data collection

|  | Dataset 1 | Dataset 2 | Dataset 3 | Dataset 4 |
| --- | --- | --- | --- | --- |
| Acceleration voltage (kV) | 200 | 200 | 200 | 300 |
| Pixel size (Å) | 1.054 | 1.054 (0.525) | 1.054 (0.525) | 1.06 (0.53) |
| Defocus range (µm) | -1.2 to -2.5 | -1.2 to -2.5 | -1.2 to -2.5 | -1.2 to -2.5 |
| Frames per movie | 40 | 40 | 40 | 40 |
| Total dose (e<sup>-</sup>/Å<sup>2</sup>) | 40 | 42 | 41.34 | 41.6 |
| Number of micrographs | 513 | 1816 | 9192 | 15 989 |
| Number of particles | 30 743 | 83 575 | 206 256 | 657 305 |
| Particles in final refinement | 14 949 | 42 390 | 114 880 | 113 721 |

### Model building, refinement and validation

A model of the *V. necatrix* ribosome (pdb 6RM3^16^) was used as a starting template to build each individual protein and RNA chain into the single particle reconstruction maps using Coot^28^. The individual maps were superimposed onto a reference whole-ribosome map using Chimera^29^ and the CCP4 software suite^30^. Individual chains were positioned into the *S. lophii* ribosome map using Chimera, Coot or the phased molecular replacement option in MOLREP^31^ adapted for use in EM^32^. The density maps were of high quality and allowed to build the full atomic model for most of the rRNA and of 71 proteins, in many cases extending the chains that were present in the *V. necatrix* template model. Cryo-EM density peaks within 2.0-2.1 Å of oxygen ligands were modelled as Mg^2+^ ions (Fig. S12B). Peaks with both oxygen and nitrogen ligands and metal-ligand distances exceeding 2.6 Å were modelled as potassium ions (Fig S12C), these were in line with potassium ion assignment by long-wavelength X-ray crystallography in the *Thermus thermophilus* ribosome^33^. The residues of the C-terminal part (77-103) of the protein eL24 that were missing in the *V. necatrix* structure (pdb 6RM3) were built as a polyalanine model. Later, a full atomic model of eL24 was predicted using Alphafold2 software^34^, which allowed sequence assignment for residues of the polyalanine model. Additionally, the Alphafold2 prediction of the C-terminal (non-containing Ub) part of the eS31 protein was successfully positioned in the small subunit head density map with addition of Zn^2+^ ion (Fig. S13). Each protein model was rebuilt and validated using the ISOLDE software^35^, implemented in ChimeraX^36^. The model was refined using REFMAC5^37^ from the CCPEM package^38^. The structure of *S. lophii* uL1 was predicted with Alphafold2 and the position of it obtained within the L1 stalk map by aligning 5T62 from *S. cerevisiae*^39^. The deacylated tRNA in the E-site was obtained by overlaying structure 6H4N from *E. coli* as a starting reference^9^. Two *S. lophii* ribosome particles were fitted into the sub-tomogram average using ChimeraX^36^, and the L1 stalk, P-stalk and MDF1 modelled into the additional density observed in the dimer map, using Alphafold2 to model individual proteins.

### Nano-LC Mass Spectrometry

Samples were run on a 10% SDS-PAGE gel until the dye front had migrated approximately 1 cm into the separating gel. Each gel lane was then excised as a single slice and subjected to in-gel tryptic digestion using a DigestPro automated digestion unit (Intavis Ltd.).

The resulting peptides were fractionated using an Ultimate 3000 nano-LC system in line with an Orbitrap Fusion Tribrid mass spectrometer (Thermo Scientific). In brief, peptides in 1 % (vol/vol) formic acid were injected onto an Acclaim PepMap C18 nano-trap column (Thermo Scientific). After washing with 0.5 % (vol/vol) acetonitrile 0.1 % (vol/vol) formic acid, peptides were resolved on a 250 mm × 75 μm Acclaim PepMap C18 reverse phase analytical column (Thermo Scientific) over a 150 min organic gradient, using 7 gradient segments (1-6 % solvent B over 1min., 6-15 % B over 58 min, 15-32 % B over 58 min, 32-40 % B over 5 min, 40-90 % B over 1 min, held at 90 % B for 6 min and then reduced to 1 % B over 1min.) with a flow rate of 300 nl min^−1.^ Solvent A was 0.1 % formic acid and Solvent B was aqueous 80 % acetonitrile in 0.1 % formic acid. Peptides were ionized by nano-electrospray ionization at 2.2 kV using a stainless-steel emitter with an internal diameter of 30 μm (Thermo Scientific) and a capillary temperature of 275 °C.

All spectra were acquired using an Orbitrap Fusion Tribrid mass spectrometer controlled by Xcalibur 2.1 software (Thermo Scientific) and operated in data-dependent acquisition mode. FTMS1 spectra were collected at a resolution of 120 000 over a scan range (m/z) of 350-1550, with an automatic gain control (AGC) target of 400 000 and a max injection time of 100 ms. Precursors were filtered according to charge state (to include charge states 2-7), with monoisotopic peak determination set to peptide and using an intensity range from 5E3 to 1E20. Previously interrogated precursors were excluded using a dynamic window (40 s +/-10 ppm). The MS2 precursors were isolated with a quadrupole mass filter set to a width of 1.6m/z. ITMS2 spectra were collected with an AGC target of 5000, max injection time of 50ms and HCD collision energy of 35 %.

The raw data files were processed and quantified using Proteome Discoverer software v2.1 (Thermo Scientific) and searched against the UniProt *S. lophii* [1358809] database (downloaded November 2020; 2499 sequences) using the SEQUEST HT algorithm. Peptide precursor mass tolerance was set at 10 ppm, and MS/MS tolerance was set at 0.6 Da. Search criteria included oxidation of methionine (+15.995 Da), acetylation of the protein N-terminus (+42.011Da) and methionine loss plus acetylation of the protein N-terminus (-89.03 Da) as variable modifications and carbamidomethylation of cysteine (+57.021 Da) as a fixed modification. Searches were performed with full tryptic digestion and a maximum of 2 missed cleavages were allowed. The reverse database search option was enabled and all data was filtered to satisfy false discovery rate (FDR) of 5 %. Raw data files were also searched against a hypothetical translation of the Celtic Deep (GCA_001887945.1, translated by getorf, min 60 nucleotides) to identify any short hypothetical genes not annotated in the *S. lophii* genome project.

### Sequence analysis

Ribosomal RNA genes were retrieved from whole genome shotgun sequence and TSA records (MQSS01000307.1 corrected with TSA sequence GALE01012333, MQSS01000329.1 and GALE01012333.1). Ribosomal protein sequences were retrieved from the *Spraguea lophii* genome Bioproject AMN02141961^40^. Not all ribosomal proteins were annotated in this original genome sequence due to the fact that the assembly was fragmented, some shorter ribosomal proteins were missed in the annotation, and the fact that some of them have short, rarely spliced introns that interrupt the ORFs^41^. To find the full set of ribosomal proteins in *S. lophii*, the assemblies of 10 environmental samples of *Spraguea* were searched using tblastn^42^. For poorly conserved proteins such as MDF2, msL1 and Lso2, that were not retrieved by this approach, PSI-BLAST and grep searches of conserved motifs in a predicted translation of the assembly in all 3 frames was used to look for evidence of their presence in the *Spraguea* genomes.

### Data deposition

The atomic coordinates of the *S. lophii* ribosome monomer and the single particle reconstruction density maps were deposited to the Protein Data Bank (https://www.rcsb.org) with accession number 7QCA and to the Electron Microscopy Data Bank (https://www.ebi.ac.uk/emdb) with the accession number EMD-13892. The docked coordinates and density map of the *S. lophii* L1 stalk were deposited to the Protein Data Bank with accession number 7YZL and EMD-14390. The sub-tomogram average of the *S. lophii* ribosome dimer and the coordinates of two ribosome particles fitted into it were deposited to the Protein Data Bank with accession number 7QJH and EMD-14020.

## Results

### *S. lophii* ribosomes form translationally dormant 100 S dimers prior to host infection

To investigate the structure of hibernating ribosomes *in situ*, we germinated microsporidian spores of the species *S. lophii* on the cryoEM grid and plunge-froze these grids in liquid ethane. Examination of the samples in the electron microscope showed spores that were too dense to be penetrated by the electron beam. Many of these spores exhibited up to 50 µm long and 150 – 200 nm wide extensions, which were identified as polar tubes (PTs); Fig. S1). We recorded cryoEM projections, as well as tomograms of PT segments and analysed the data in detail. Close inspection of the PTs revealed that they were confined by a fuzzy, likely proteinaceous coat, which was often lined with a membrane on the inside (Figs. S1H and S2). These membrane-lined PTs contained clearly recognizable cytosolic content, within which ribosomes could be distinguished (Figs. 1A, S1H and S2).

**Figure 1.**
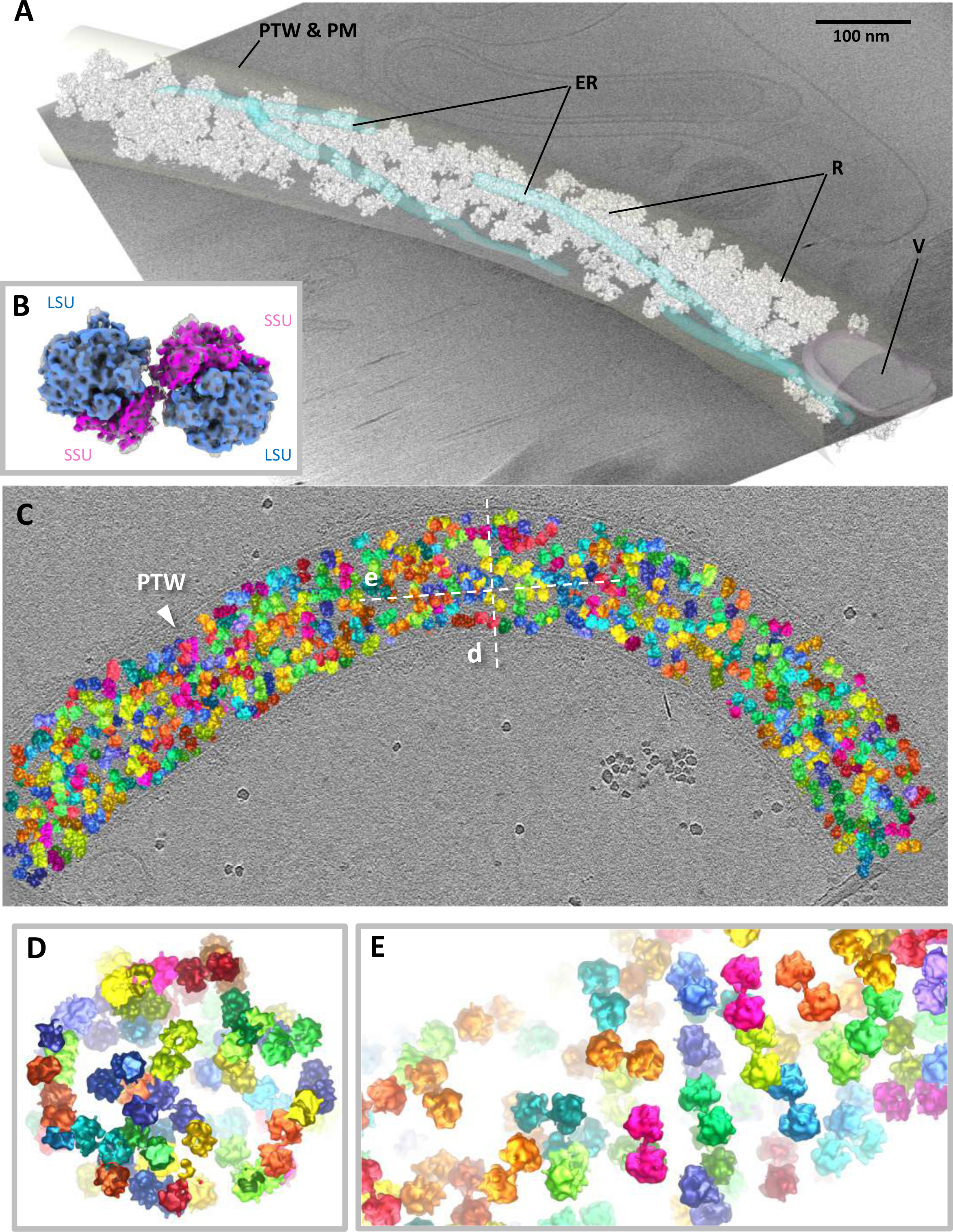
Structure and organisation of the *S. lophii* ribosome dimer *in situ*. **A** Segmented tomogram of a polar tube (PT), showing endoplasmic reticulum (ER, transparent blue), a vesicle (V, transparent magenta) and ribosomes (R, white). The polar tube wall (PTW) and plasma membrane (PM) are transparent grey. **B** Sub-tomogram average of the *S. lophii* ribosome dimer at 10 Å resolution, showing the small subunit in pink and the large subunit in blue. **C** Organization of ribosome dimers inside a PT (various colours). **D** and **E** Cross sections through the PT from areas indicated by the dotted lines (d and e) in C.

5001 ribosomes within the polar tubes from *S. lophii* were selected manually and subjected to sub-tomogram averaging in Relion 3.1 followed by emClarity (Fig. S3). The resulting map (Fig. 1B, Fig. S4) at a resolution of 10.3 Å (Fig. S5) shows a dimeric complex of two 70 S ribosomes interacting via the small subunit in an antiparallel head-to-head orientation. 3D classification indicated that the majority of the ribosomes in the PT exist as dimers. Only 1.6 % (79 of the 5001 particles) contributed to a class containing density accounting for monomeric ribosomes (Fig. S6).

Placing the sub-tomogram average back into the original particle positions within the tomogram showed variable packing of ribosome dimers within the PT (Fig. 1C-E). While in some instances the ribosome dimers are almost at touching distance, other areas contain pockets of cytosol between them (Fig. 1C-E). A statistical analysis of distances and angular orientations between closest neighbours revealed no higher order of organisation (Fig. S7), indicating an absence of polysomes or any specific packing of the hibernating dimers. In addition, 3D classification also did not show any ribosomes associated with a membrane (Fig. S6), even though ER-tubules were present in some of the tomograms (Fig. 1A). These results clearly indicate that none of the ribosomes in the PTs are actively translating or engaged with the SEC translocon.

### Single particle cryoEM of hibernating *S. lophii* ribosomes

To obtain a better understanding of the structure of the hibernating ribosome from *S. lophii*, we isolated the ribosomes from spores using sucrose gradient centrifugation, prepared cryoEM samples using graphene oxide grids and recorded single particle datasets. 2D classification showed that the particles were 70 S ribosome monomers (Fig. S8), suggesting that the sample preparation procedure did not maintain the dimer contacts. This is in accordance with the single particle structures of isolated hibernating ribosomes of the microsporidia *V. necatrix, P. locustae and E. cuniculi,* where dimers were also not observed.

Using multibody refinement in Relion 3.1, we were able to obtain a map of the *S. lop*hii 70 S ribosome at 2.26 – 2.78 Å resolution (Fig. S9, S10 and S11), which exceeds the resolution previously achieved for microsporidian ribosomes^16–18^. Based on this map, we built an atomic model of the *S. lophii* ribosome, consisting of 71 protein chains and 96.8% of the rRNA sequence into our density (Fig. S12 and S13). Comparing our structure with previously published data showed that the hibernating ribosome of *S. lophii* is in its non-rotated conformation (Fig. S14), similar to structures of the hibernating ribosome of *P. locustae* but in contrast to those of *V. necatrix* and *E. cuniculi.* As is seen in the other structures, the *S. lophii* ribosome shows a drastic reduction in the expansion segments of the ribosomal RNAs relative to other eukaryotes, however as in *P. locustae,* predicted 18 S and 28 S rRNA are slightly longer than counterparts in *E. cuniculi* (68 and 129 nucleotides longer respectively), showing that rRNA reduction is slightly less severe in these earlier branching lineages.

We did not observe a density for the large ribosomal protein subunit eL29, which was present in the previously published structures of microsporidian ribosomes from *V. necatrix, P. locustae* and *E. cuniculi.* In accordance with this, we find that the corresponding gene is absent from the *S. lophii* genome, suggesting a loss within this microsporidian lineage. However, our map does contain a density for the C-terminal domain of the small subunit protein eS31 (one of two ribosomal ubiquitin fusion proteins; structure predicted by Alphafold2, Figure S15). This subunit was also observed in the *V. necatrix and E. cuniculi* ribosomes but is missing from the *P. locustae* structure.

Interestingly, in our monomeric ribosome structure we did not observe densities that could account for the hibernation factors seen in the previously determined single particle structures of microsporidian ribosomes from *V. necatrix, P. locustae* and *E. cuniculi*. No densities suggesting the presence of MDF1^16, 18^ or Lso2^17^ were evident in the A,P or E sites. In addition, the polypeptide exit tunnel did not contain a blocking hibernation factor, such as MDF2 or Lso2 (Fig S16). Multiple rounds of 3D classification did not reveal any variability in this region.

Based on these observations, we investigated which hibernation factors are present in and transcribed from the genome of *S. lophii*. Searches of the *S. lophii* genomes using BLASTP and TBLASTN for known microsporidian hibernation factors showed that homologues of MDF1 and Lso2 are encoded in the genome while MDF2 could not be detected (Fig. S17). MDF2 is a protein that has only been identified in *V. necatrix, N. ceranae* and *N. apis*, and it is possible that it is either an innovation within the *Vairimorpha* and *Nosema* lineage, or present but hard to detect in more distantly related species. Interrogating RNAseq data of *S. lophii* detected reads that could be mapped to MDF1 and Lso2, suggesting that both genes are expressed. Curiously however, read coverage did not stretch to the 5’ end of the Lso2 gene but started 17 nucleotides into the predicted gene (Fig. S17). Mass spectrometry analysis of our *S. lophii* samples revealed that MDF1 was present in some fractions of the ribosome purification, but absent in others (Fig. S18).

Instead of harbouring previously described hibernation factors, our single particle maps show a different, weak density near the ribosome’s E site (Fig. S19). Focused classification and refinement of this part of our single particle map resulted in a map with local resolution ranging between 3 and 6 Å (Fig. S11C). This map enabled us to identify this density as the L1 stalk in closed conformation, which is bound to a tRNA located in the E site of the ribosome (Fig. 2B, D, E). Further 3D classification did not reveal ribosomes that lacked the density for the E site tRNA, suggesting that the tRNA was bound to the great majority of the ribosomes in the sample, but flexible. The rRNA and protein uL1 (predicted by Alphafold2) of the L1 stalk, and a deacylated tRNA were modelled into this map (Fig. 2D, E; Fig. S20). The tRNA shows three distinct interactions with the ribosome: i) the 3’ CCA terminus on the acceptor arm interacts with protein eL42 and the 23 S rRNA ii) the anticodon arm binds the 16 S rRNA and protein uS7 and iii) the elbow of the tRNA interacts with the L1 loop of the LSU rRNA and uL1 protein which make up the L1 stalk. This is reminiscent of the structure of the hibernating ribosome dimer from *E. coli*, where a deacetylated tRNA was found in the E site^9^. However, while the tRNA was stabilized by the hibernation factor HPF in the *E. coli* ribosome dimer, a similar protein is not observed in our structure. This suggests that the interactions with proteins uL1, uS7 and eL42, and with the 16 S and 23 S rRNAs, are sufficiently stable to fix the tRNA in the E site in the absence of a hibernation factor in the *S.lophii* ribosome.

**Figure 2.**
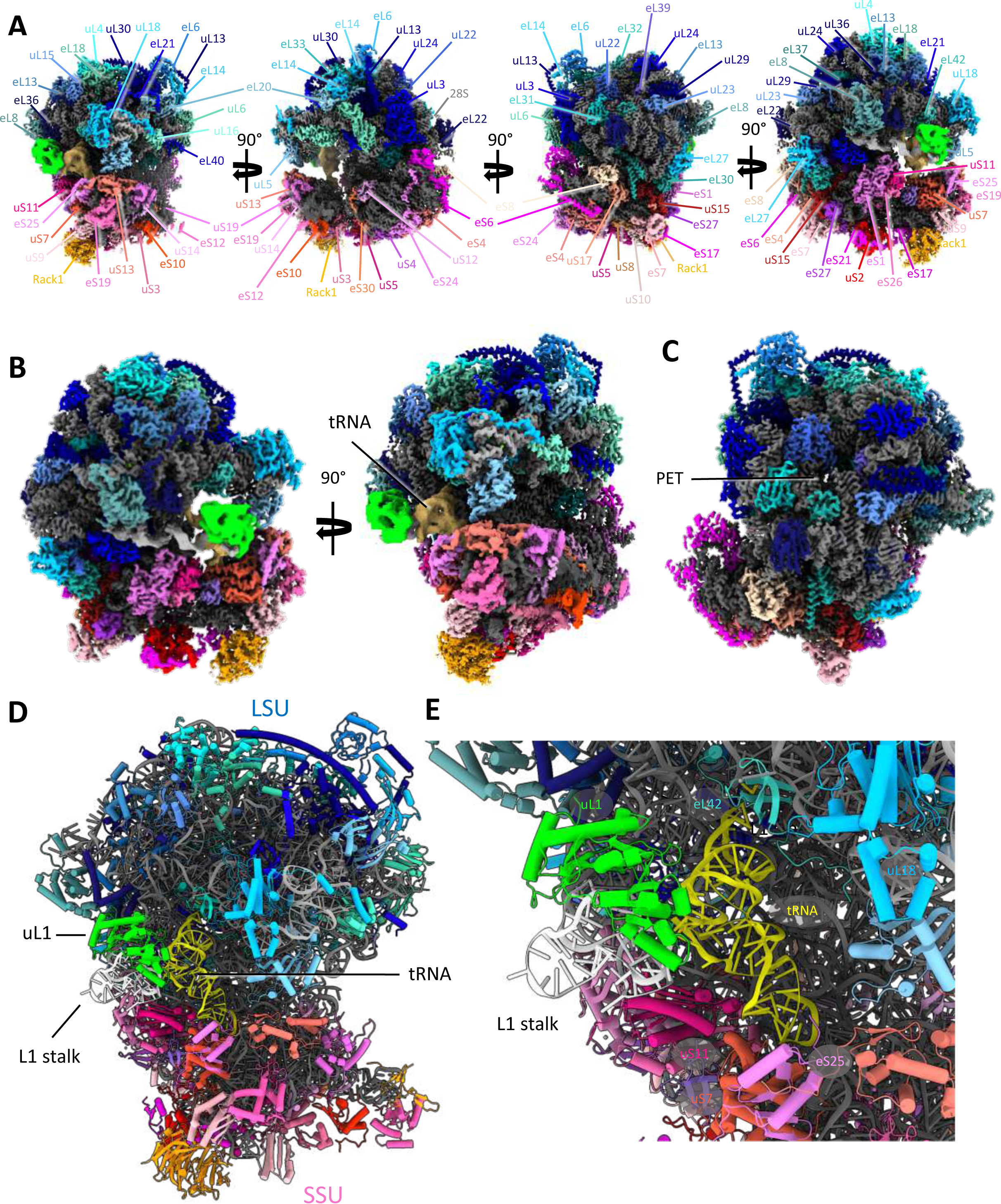
– Single particle structure of the *S. lophii* 70 S ribosome at 2.2 - 2.7 Å resolution. **A** Various views of the ribosome, showing the protein subunits of the large subunit (LSU) in shades of blue, the small subunit (SSU) in shades of red and the rRNA in grey. Subunits are indicated. **B** The L1 stalk shown in two views, rotated by 90 degrees. A tRNA (beige) is found in the E-site of the ribosome. The rRNA of the L1 stalk is shown in light grey with the uL1 subunit in green. **C** The polypeptide exit tunnel (PET) appears to be free of hibernation factors. **D** View of the E site of the ribosome. **E** Closeup of D. uL1 (green) and deacetylated tRNA (yellow) have been modelled into the density of the L1 stalk and E site. The tRNA interacts with protein uS7 of the small subunit, proteins eL42 and uL1 of the large subunit and rRNA.

### A unique ribosomal dimerization mechanism

To obtain a detailed structure of the eukaryotic hibernating ribosome dimer, we fitted our atomic model into our dimer map obtained by sub-tomogram averaging (Fig. 3A, B). This revealed a 100 S particle that adopts a conformation that is entirely different compared to its bacterial counterparts. Even though the orientations of the 70 S ribosomes within the 100 S dimers from *E. coli, S. aureus and T. thermophilus* differ by a rotation around the dimer interface^43^, the dimer contact is always established between the subunits uS2 (via RPF in *E. coli* and HPF in *S. aureus and T. thermophilus*) (Fig. 3 D, E). In contrast, the eukaryotic dimer interface observed in *S. lophii* is located between protein eS31 and the 16 S rRNA in the beak of the small subunit. A connecting bridge of density is observed between eS31 and the 16 S small subunit rRNA from one ribosome, and the identical sites on the other ribosome in the dimer (Fig. S21). This suggests an entirely different mechanism of dimer formation to those seen in bacteria. In line with this, no homologues to known bacterial dimerization factors were identified from our mass spectrometry or genomic sequence analyses.

**Figure 3.**
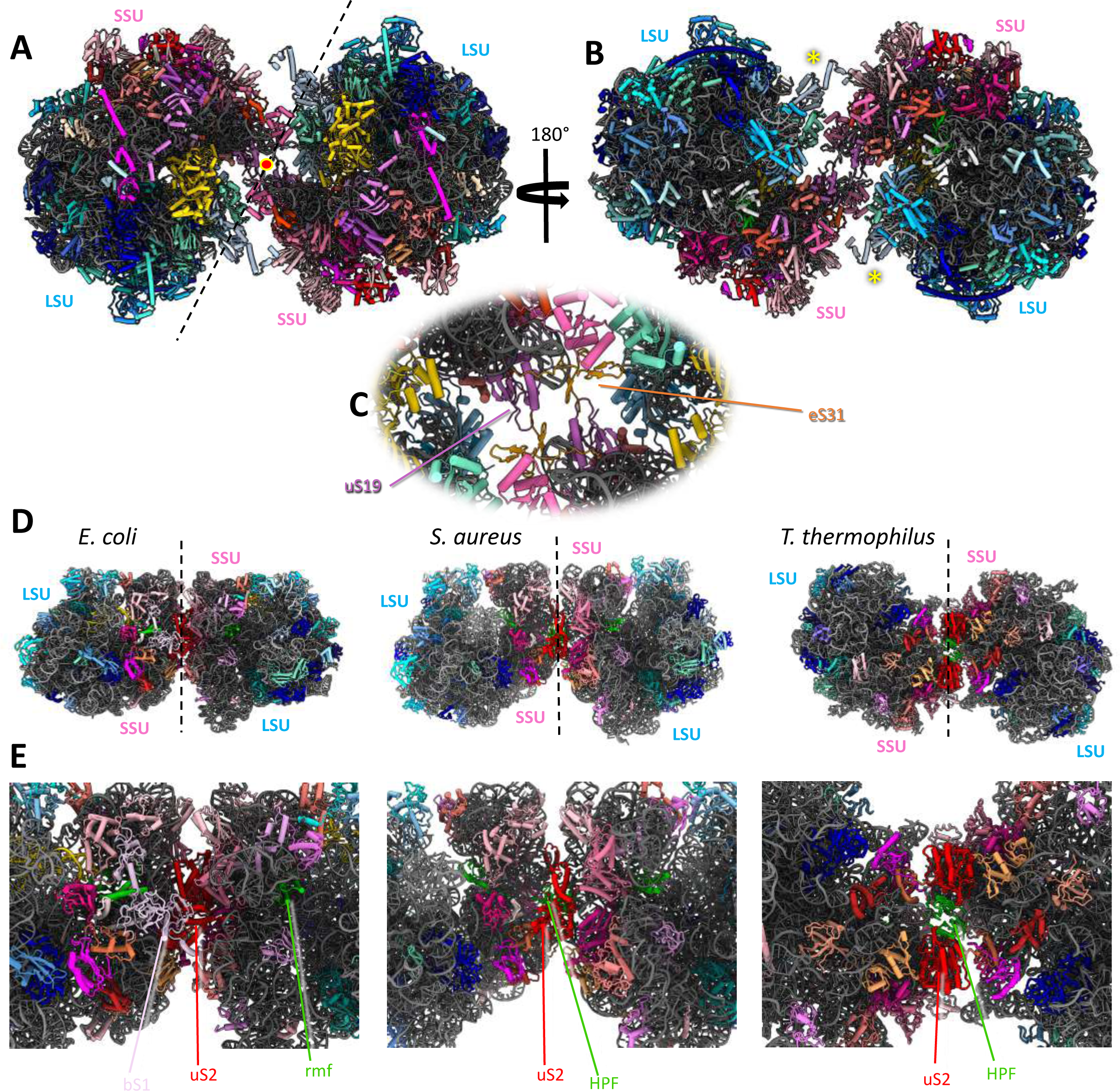
Model of the hibernating ribosome dimer from *S. lophii.* **A, B** Two views of the ribosome dimer rotated by 180° degrees. In **A**, a red dot indicates the C2 symmetry axis and a dashed line the plane of the dimer interface. Structures coloured as in Fig. 2, but additionally eukaryotic elongation factor 2 is shown in gold and MDF1 in lime **C** Closeup of the dimer interface, which involves the protein eS31 and the 16 S rRNA of the SSU. The P-stalk (yellow asterisk; panel B) is observed in a fixed position, projecting close to the dimer interface, occluding it from interacting with regulatory factors involved in translation initiation and elongation. **D** A gallery of known hibernating 100 S ribosomes from bacteria in comparison and closeups of their dimer interfaces (**E**). The orientation of the 70 S ribosomes within each of the bacterial dimers is different compared to the one of *S. lophii.* In each case, the dimer interfaces are established via the ribosomal subunit uS2 and the dimerisation factor HPF *(S. aureus* and *T. thermophilus),* and the additional factor bs1 in *E. coli*.

In contrast to our single particle map, the sub-tomogram average of the dimer did not contain a density for an E site tRNA. In contrast, this region was occupied by a density that closely matched the structure of MDF1 (Fig. S22). Fitting the Alphafold-2 prediction of *S. lophii* MDF1 into the density revealed that MDF1 mimics the position of an E site tRNA, as previously suggested^18^, holding the L1 stalk in a closed position. In fact, our data show that MDF1 is readily exchanged by an E site tRNA. When comparing our single particle structure with the sub-tomogram average, we find that the L1-stalk when MDF1-bound is closed even further than when tRNA-bound (Fig. S23). This indicates that MDF1 holds the L1 stalk in a conformation that is incompatible with tRNA binding.

Furthermore, our dimer map showed a clear density for the P-stalk, which was not resolved in the single particle structure. We were able to model the dimer-borne P-stalk based on the structure of a pig ribosome, 3J7P^44^ and Alphafold models of the individual *S. lophii* components (Fig S24). In eukaryotes this region contains the ribosomal subunits uL10, uL11 and P1/P2 with uL10 forming the base of the stalk to which a number of dimers of P1/P2 can bind, and we were able to model proteins uL10 and uL11 into the P-stalk density. This stalk region is important for the binding of translational GTPases, which catalyse various steps of translation^45^. Interestingly, we also discovered a density matching the Eukaryotic Elongation Factor 2 (EF2), the presence of which was confirmed by our mass spectrometry data and hence also modelled an Alphafold prediction of *S. lophii* EF2 into the P-stalk region (Fig S24 C). In the 100 S hibernating ribosome, the P-stalk is located close to the dimer interface where it may form interactions with proteins eS31 and eS12 from the opposite subunit of the dimer and thus stabilize the 100 S complex.

## Discussion

Here we report the *in situ* structure of a eukaryotic hibernating ribosome dimer from PTs of the microsporidian species *S. lophii*. The architecture of the dimer, as well as the dimer interface are markedly distinct when compared to bacterial hibernating 100 S ribosomes. This suggests different dimerization mechanisms in bacterial and eukaryotic 100 S ribosomes. The observation that ribosomal dimerization is retained in the otherwise strongly reduced microsporidian ribosome highlights its importance for ribosomal protection during phases of cellular dormancy. An open question to be answered is how widespread and variable hibernating ribosome dimers are amongst eukaryotes. Interestingly, one study demonstrated that hibernating ribosome dimers occur in rat glioma cells under amino acid starvation conditions^12^. This suggests the intriguing possibility that ribosomal dimerisation is conserved throughout the eukaryotic kingdom, including mammals, with unforeseen important implications for cellular homeostasis and health.

The dimer interface involves the ribosomal subunit eS31. This protein is one of two eukaryotic-specific ubiquitin fusion proteins found within the ribosome, the second one being eL40, which is situated at the base of the P-stalk. These two ubiquitin fusion proteins are universally encoded in the genomes of model eukaryotes^46^. Ubiquitin is a reversible post-translational modifier and is involved in many different cellular processes; being added to, and removed from, proteins and consequently altering their localisation or activity. In the ribosome, ubiquitin domains are themselves proteolytically removed from both eS31 and eL40^47^. A recent review questions whether the selective advantage of ubiquitin being fused to eS31 and eL40 could be a means of coupling the synthesis and degradation of proteins to maintain proteostasis in eukaryotes. It was also discussed that the specific fusions of ubiquitin with eS31 and eL40 over any other ribosomal proteins suggest an important role for ubiquitin in eS31 and eL40 expression, ribosome assembly or ribosome function^46^. Our observation that both eS31 and eL40 are involved in or close to the dimer interface may thus point to a role in ubiquitin processing of 100 S ribosome formation in eukaryotes.

Moreover, eS31 and eL40 both contain a highly conserved zinc-binding motif (totally conserved in the top 100 blast hits), and we have modelled zinc into our single particle structure (Fig. S13D, E). Ribosome hibernation has been shown to occur in *Mycobacterium smegmatis* as a direct result of zinc starvation, with Mycobacteria and other bacteria remodeling their ribosomes in response to zinc depletion by replacing zinc-binding ribosomal proteins with zinc-free paralogues, and releasing zinc for other metabolic processes^48, 49^. The authors proposed that ribosome hibernation is a specific and conserved response to zinc depletion in mycobacteria. The presence of the highly-conserved zinc-binding eS31 protein at the dimerisation interface raises the intriguing potential of zinc-binding playing a role in eukaryotic ribosome dimerisation in response to stress. Taken together, eS31 and possibly eL40 may be important signaling hubs for the initiation and termination of ribosomal hibernation and mediators for the formation of 100 S ribosomes.

The P-stalk has previously been shown to increase the local concentration of the translational guanosine exchange factors EF1A and EF2 while flexibly moving on the ribosome; hence promoting polypeptide elongation^50^. In the hibernating ribosome dimer, the P-stalk is orientated towards and likely participates in the dimer interface. This appears to lock the usually flexible P-stalks into a fixed position – evidenced by the fact that it was resolved in our dimer structure, as opposed to most single particle structures of individual ribosomes. It is likely that the static position of the P-stalk would reduce its ability to exchange EF1A and EF2, resulting in the halting of translation.

Previous studies of purified hibernating ribosomes from the microsporidian species *P. locustae, V. necatrix and E. cuniculi*^16–18^ did not reveal dimers. This suggests that either ribosome dimers do not exist in those species or that, consistent with our single particle data, the dimer interface is fragile and easily disrupted during sample preparation. Our data reveal the presence of the hibernation factor MDF1 in the ribosome dimer, which is exchanged by a likely deacetylated E site tRNA in the purified ribosome. MDF1 appears to cause a more complete closure of the L1 stalk when compared to the monomeric structure with deacylated tRNA in the E site. In these conditions, the ribosome is rendered inaccessible for mRNA and A and P site tRNAs. In contrast, neither our ribosome dimer structure, nor that of the monomer reveal the presence of a factor that blocks the polypeptide exit tunnel, as it is the case for *P. locustae* and *V. necatrix.* This is consistent with the absence of MDF2 in the *S.lophii* genome and suggests that blocking the polypeptide exit tunnel is not an essential requirement for ribosome hibernation, at least in some species.

Isolating ribosomes from microsporidian spores resulted in a homogenous suspension of fully assembled ribosome monomers. This suggests that the dormant dimers are readily transformable to translationally competent monomers. The presence of an E site tRNA in our structure of the purified 70 S ribosome is akin to the deacetylated E site tRNAs found in the hibernating ribosomes from *E. coli*^9^. It has been demonstrated in bacteria that such deacetylated tRNAs bind to ribosomes under conditions of amino acid shortage and tag ribosomes for dimerisation^4^. Microsporidia shut down their metabolism during spore formation in a process that likely is also accompanied by a scarcity of amino acids^51^. It is thus possible that binding of deacetylated tRNAs also plays a role in the onset of ribosomal hibernation and dimerisation in microsporidia.

Taken together, we show that the microsporidian sporoplasm is densely packed with ribosome dimers that are dormant but otherwise completely assembled and translationally competent. This shows that microsporidian spores maintain their ribosomes in an inactive, yet primed state, ready to reactivate once a host cell has been infected. Once the microsporidian parasite has invaded a host cell, the 100 S ribosome dimers must convert into 70 S monomers and shed their hibernation factor (MDF1) to become fully translationally active. In principle, this may simply ensue through mechanical disruption of the dimer interface, as observed during our single particle sample preparation, or due to a concentration effect as observed in *E. coli*, where a reduction in ribosome concentration leads to a dissociation of the dimer to monomeric ribosomes^52^. It could be envisaged that a dilution of microsporidian ribosomes takes place when the microsporidian sporoplasm leaves the densely-packed environment of the polar tube and enters the host cell. However, it is also likely that a so-far unknown microsporidian or host-cell signal is required to activate the hibernating microsporidian ribosomes. Further studies will need to be undertaken to investigate the exact sequence of events that take place during eukaryotic ribosomal hibernation and activation in microsporidia and beyond.

## Acknowledgements

LG and MM were funded by a ERC starting grant (from the European Research Council under the European Union’s Horizon 2020 research and innovation programme, grant agreement No 803894) awarded to BD. MM was also funded by a BBSRC New Investigator Research Grant (BB/R008639/1) to VG. BC and PD were funded by Wellcome Trust Seed Award (212439/Z/18/Z) awarded to BD. SRC was supported a Ministerio de Economía Y Competitividad Grant (MINECO; CAQ20"1782222-R). We acknowledge Diamond Light Source for access and support of the cryo-EM facilities at the UK’s national Electron Bio-imaging Centre (eBIC) at Diamond Light Source (under proposals BI25452 and EM18258), funded by the Wellcome Trust, MRC and BBRSC. We acknowledge access and support of the GW4 Facility for High-Resolution Electron Cryo-Microscopy, funded by the Wellcome Trust (202904/Z/16/Z and 206181/Z/17/Z) and BBSRC (BB/R000484/1). We are grateful to Ufuk Borucu of the GW4 Regional Facility for High-Resolution Electron Cryo-Microscopy for help with screening and Kate Heesom of University of Bristol Proteomics Facility for the mass spectrometry analysis.

## Figure legends

**Figure S1.**
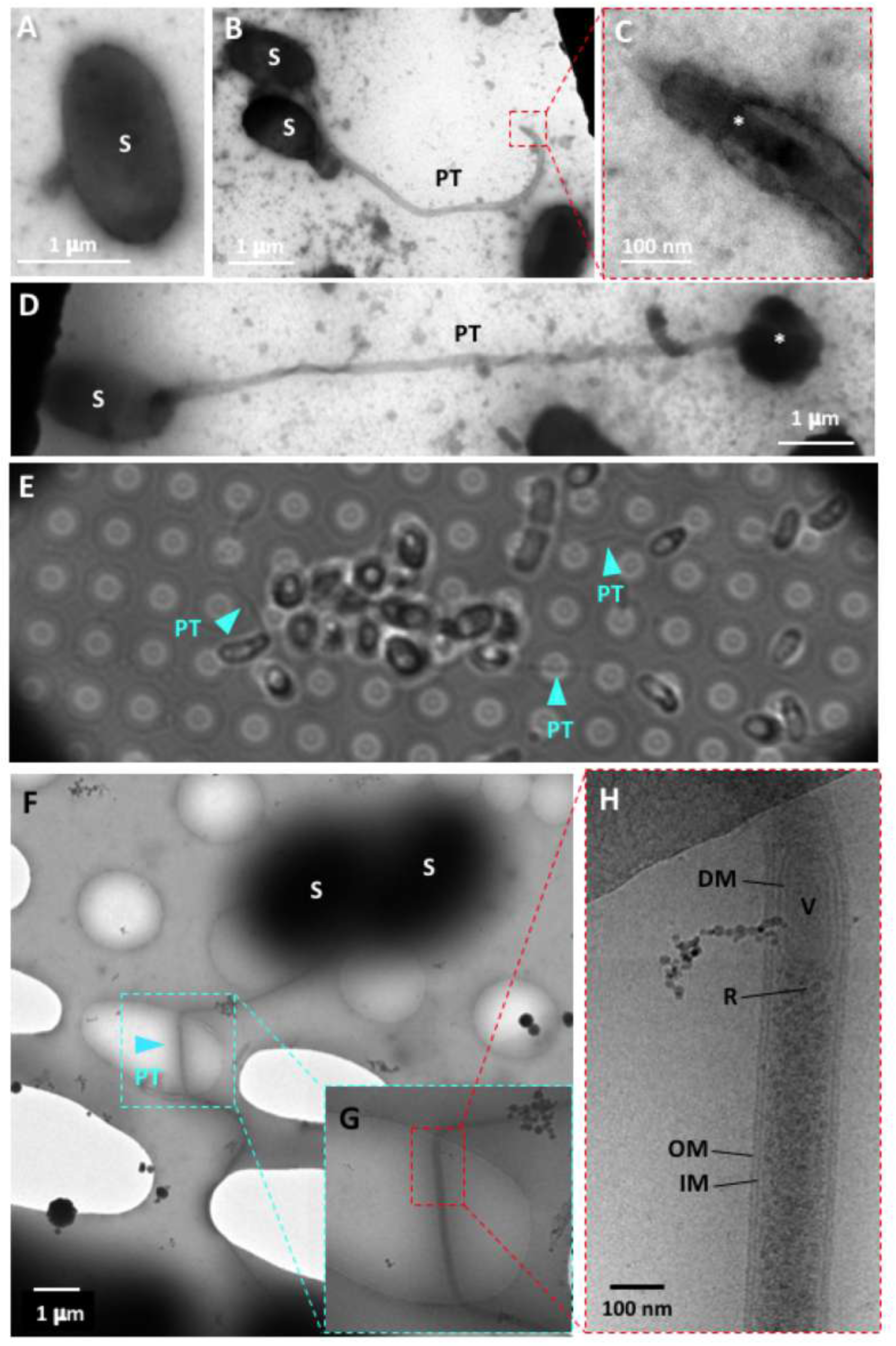
Electron microscopy of microsporidia germinated on EM grids. **A – D** Negative stain EM of **A** a dormant spore (S), **B** a germinated spore with polar tube (PT) extended, **C** the tip of a PT with sporoplasm (*) passing through at high magnification and **D** a germinated spore post transfer, with sporoplasm egressing terminally (*). **E** Bright field light microscopy showing that microsporidia can be germinated on holy carbon (Quantifoil) grids. **F-H** CryoEM of microsporidia germinated on Multi-A Quantifoil grids showing a PT emerging from a spore at three different magnifications. The PTs span Quantifoil holes, which is an important requirement for cryoET. **G**, **H** Higher magnification images of dashed area in F. **G** The PT is confined by an inner and outer membrane (IM, OM). Transported content is visible. (R) ribosomes, (V) vesicle and (DM) double membrane vesicle.

**Figure S2.**
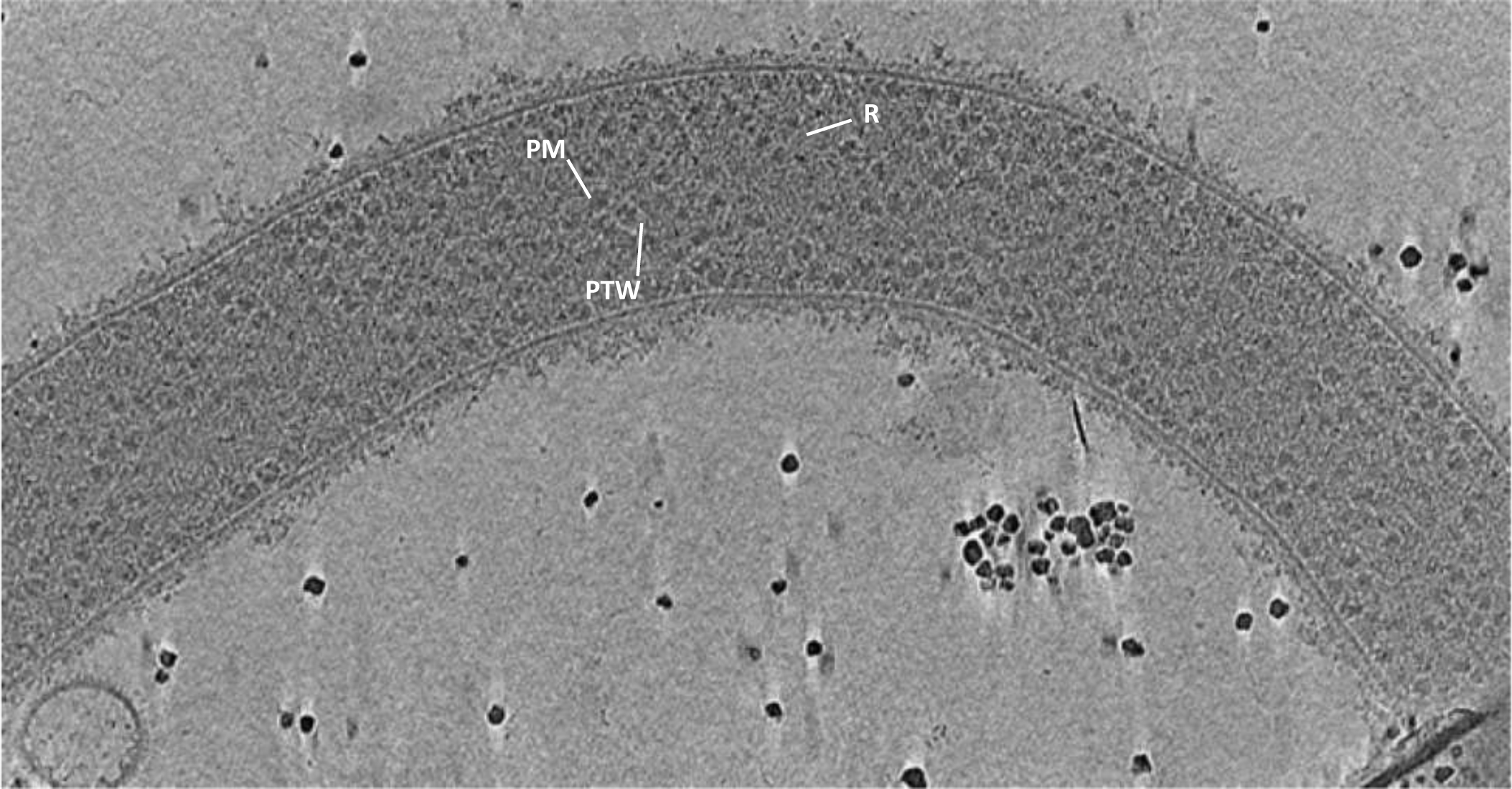
Slice through a tomogram of a polar tube of a germinated *S. lophii* spore. PTW, polar tube wall, PM, plasma membrane, R, ribosome.

**Figure S3.**
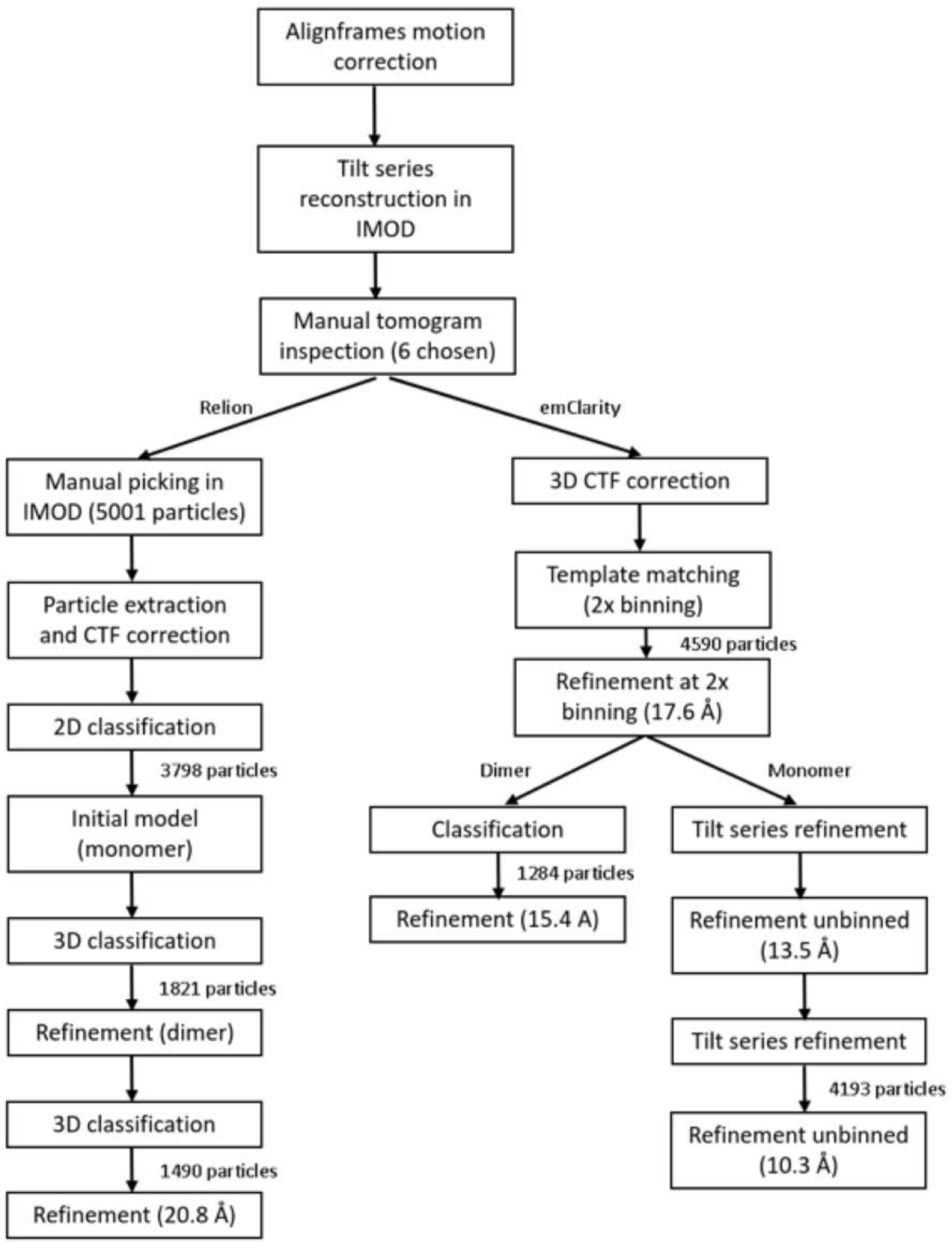
Flow chart of Relion sub-tomogram averaging processing.

**Figure S4.**
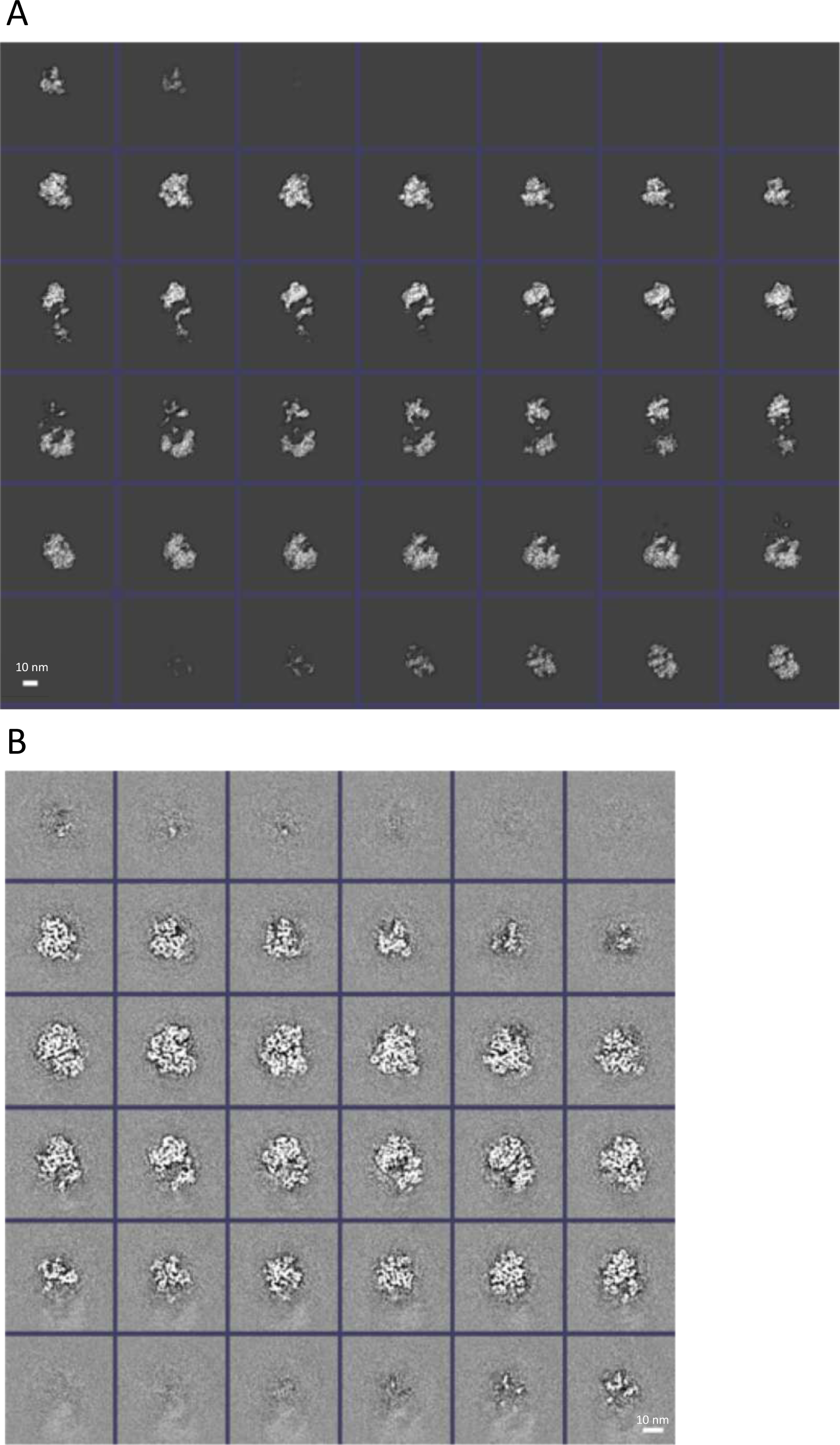
Consecutive sections through the sub-tomogram average of a ribosome dimer from *S. lophii*. **A** Average of the ribosome dimer at 15 Å resolution. **B** Average focused on one 70 S ribosome within the dimer at 10 Å resolution.

**Figure S5.**
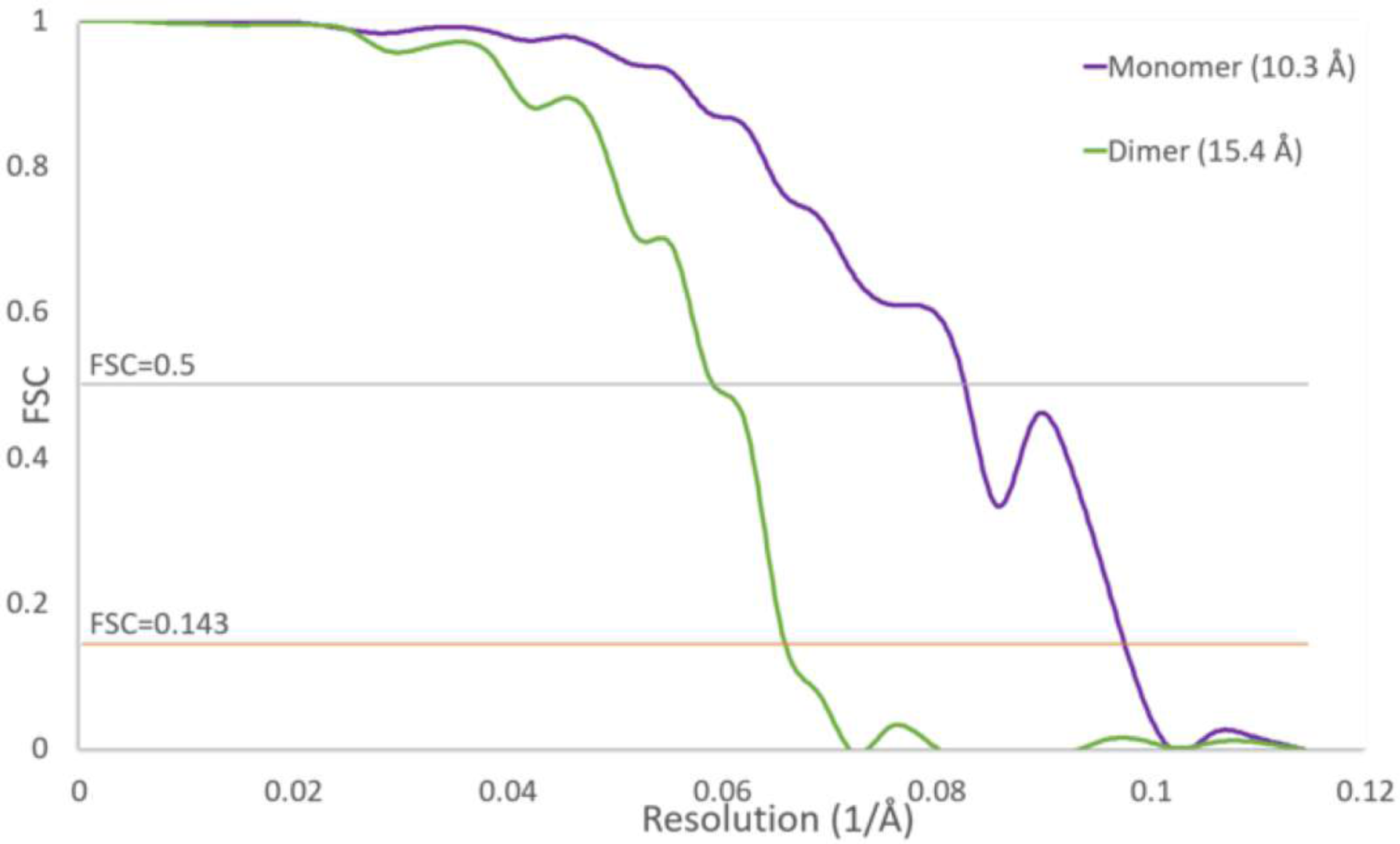
Resolution estimation of the sub-tomogram average of the *S. lophii* ribosome dimer. Gold Standard FSC of the dimer (green) and masked monomers within the average (purple).

**Figure S6.**
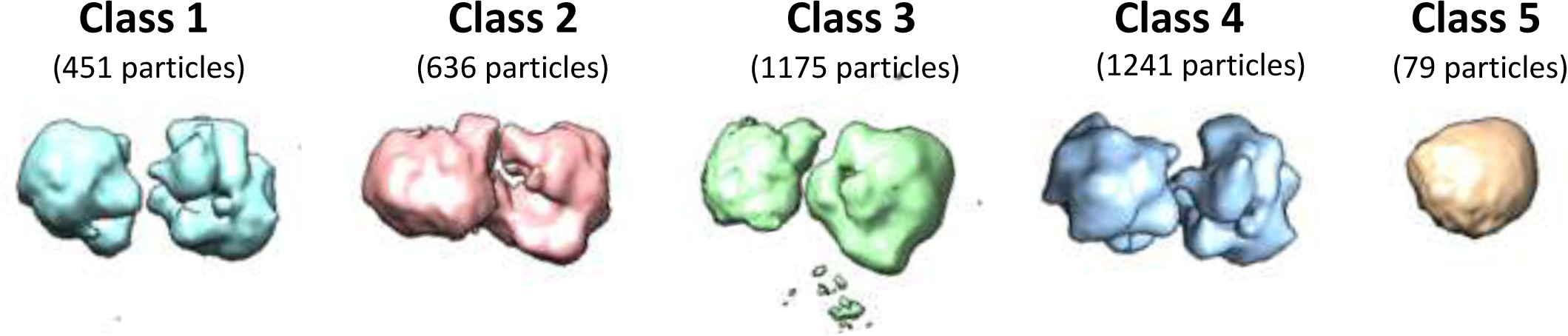
3D classification of sub-tomogram average. Out of a total of 5001 particles, only 79 particles contribute to a monomeric class. In all 100 S dimers, the 70 S ribosomes adopt the same orientation with respect to each other.

**Figure S7.**
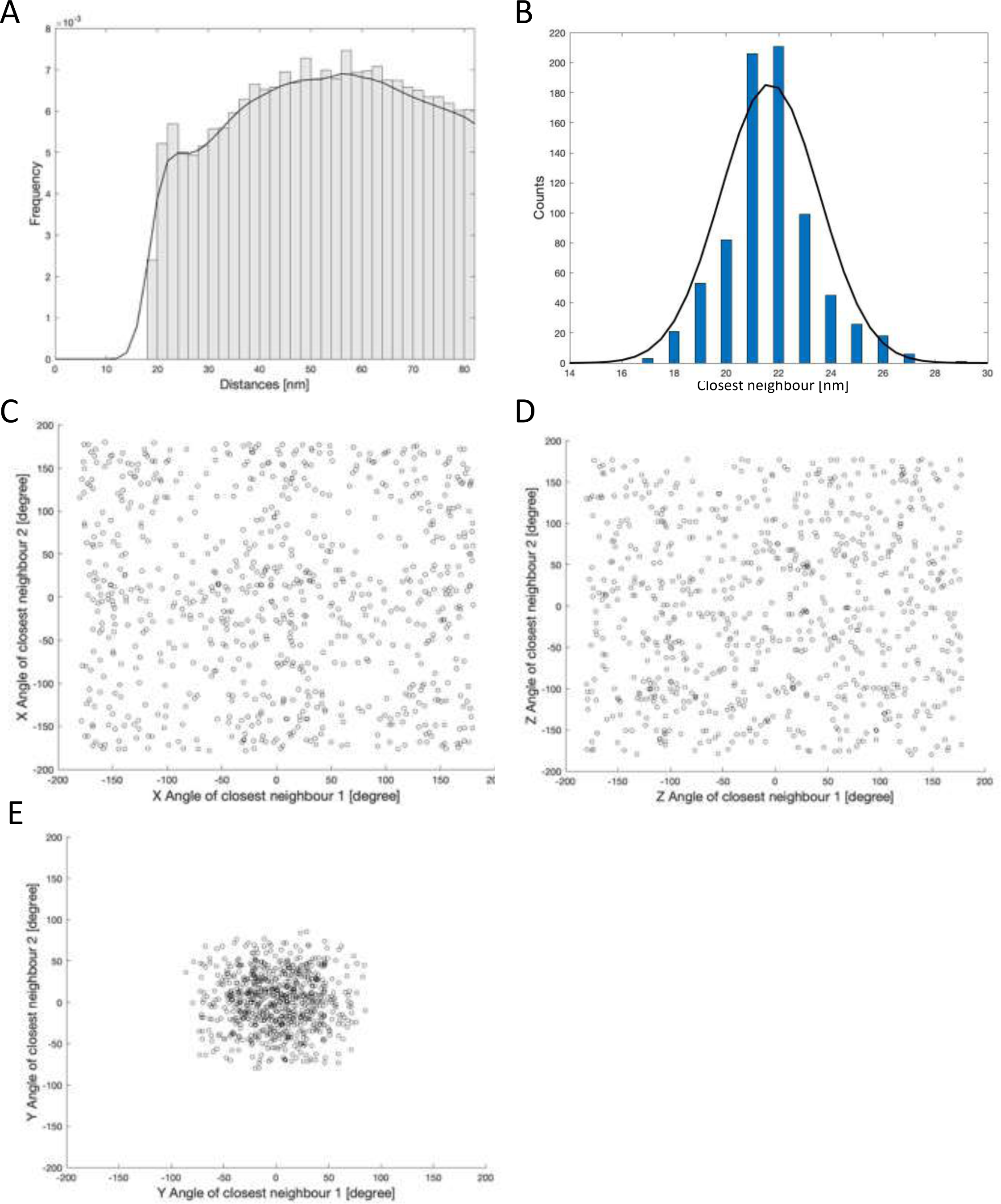
Statistical analysis of ribosome distribution within the polar tube. **A** Histogram of distances between 70 S particles in a PT with a peak at around 21 nm, corresponding to the centre-to-centre distance between two partners in a dimer pair, or ribosomes in touching distance. No other peaks or multiples of the average distance between partners that could account for the presence of polysomes are evident. **B** Histogram showing only the distribution of distances between closest neighbours, with a peak at around 21 nm. **C-E** Angular relationships between closest 70 S neighbours of all particles. Scatter plots of the X and Z angles do not show any correlation, and are randomly distributed between -180 and +180°. The Y angles of closest neighbours (i.e. dimers) of all particles range only between -90 and +90°, which is likely a manifestation of the fixed angle between two 70 S particles within a dimer.

**Figure S8.**
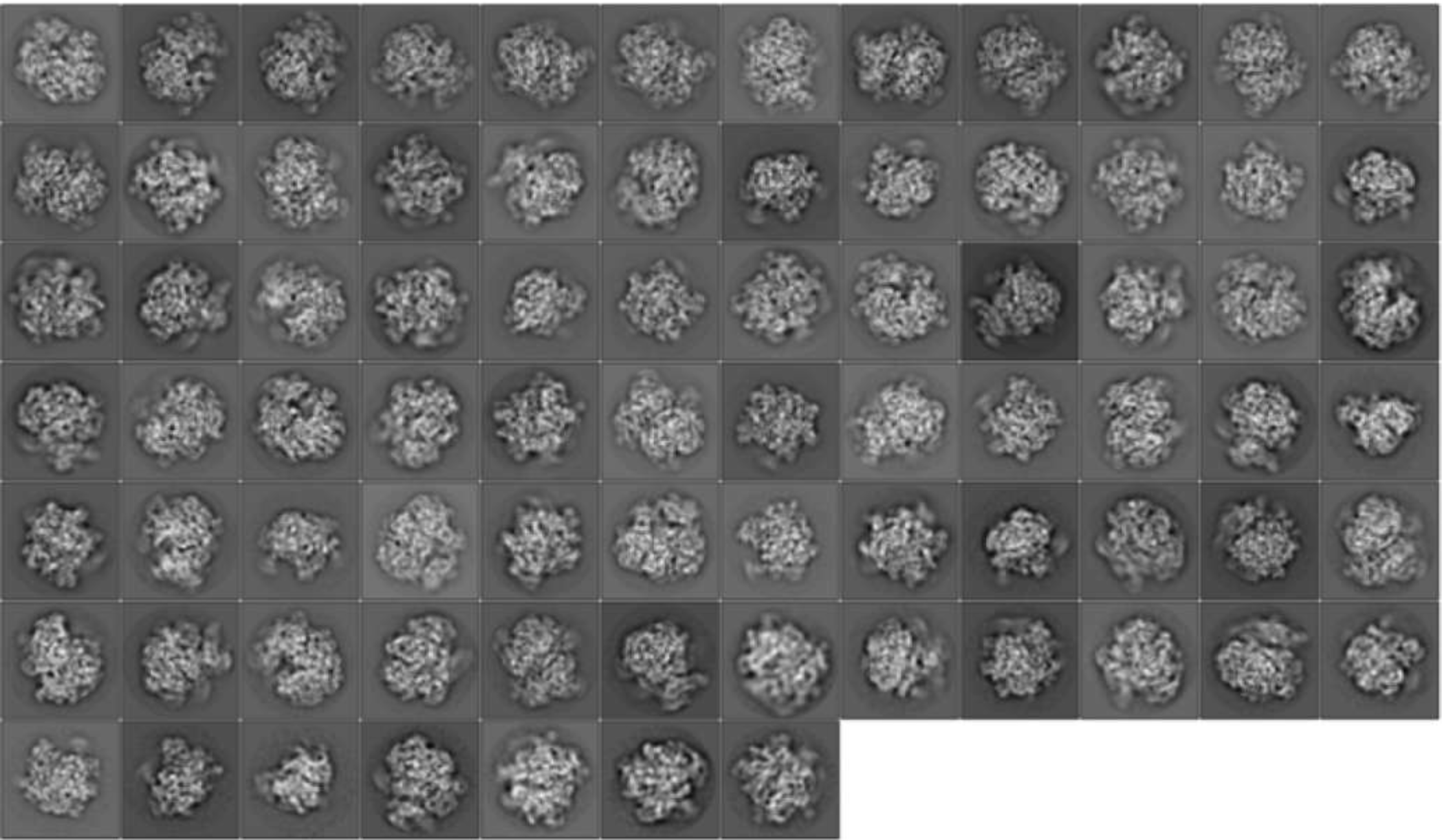
2D classification of cleaned and polished particle dataset.

**Figure S9.**
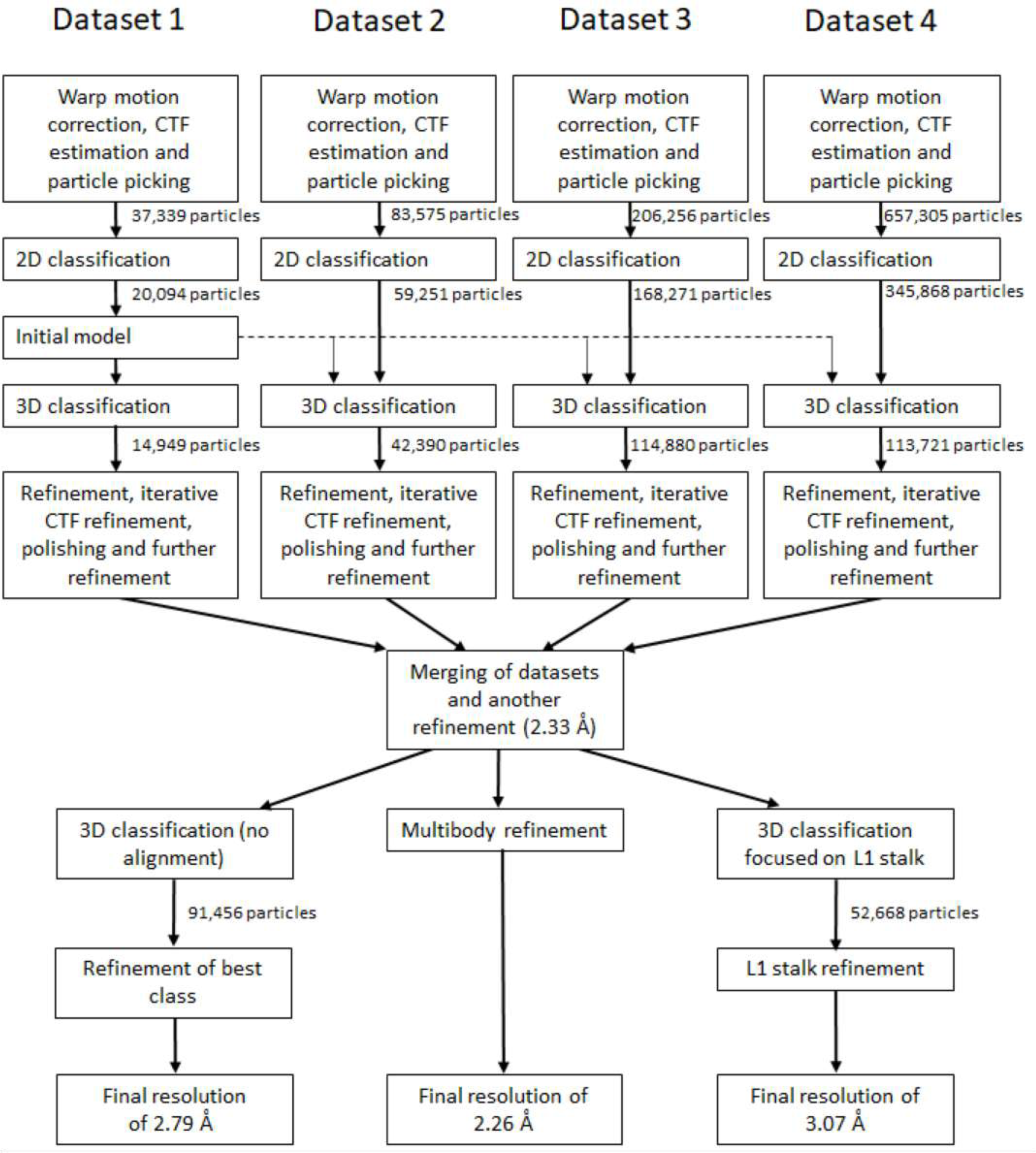
Flow chart of Relion single particle processing.

**Figure S10.**
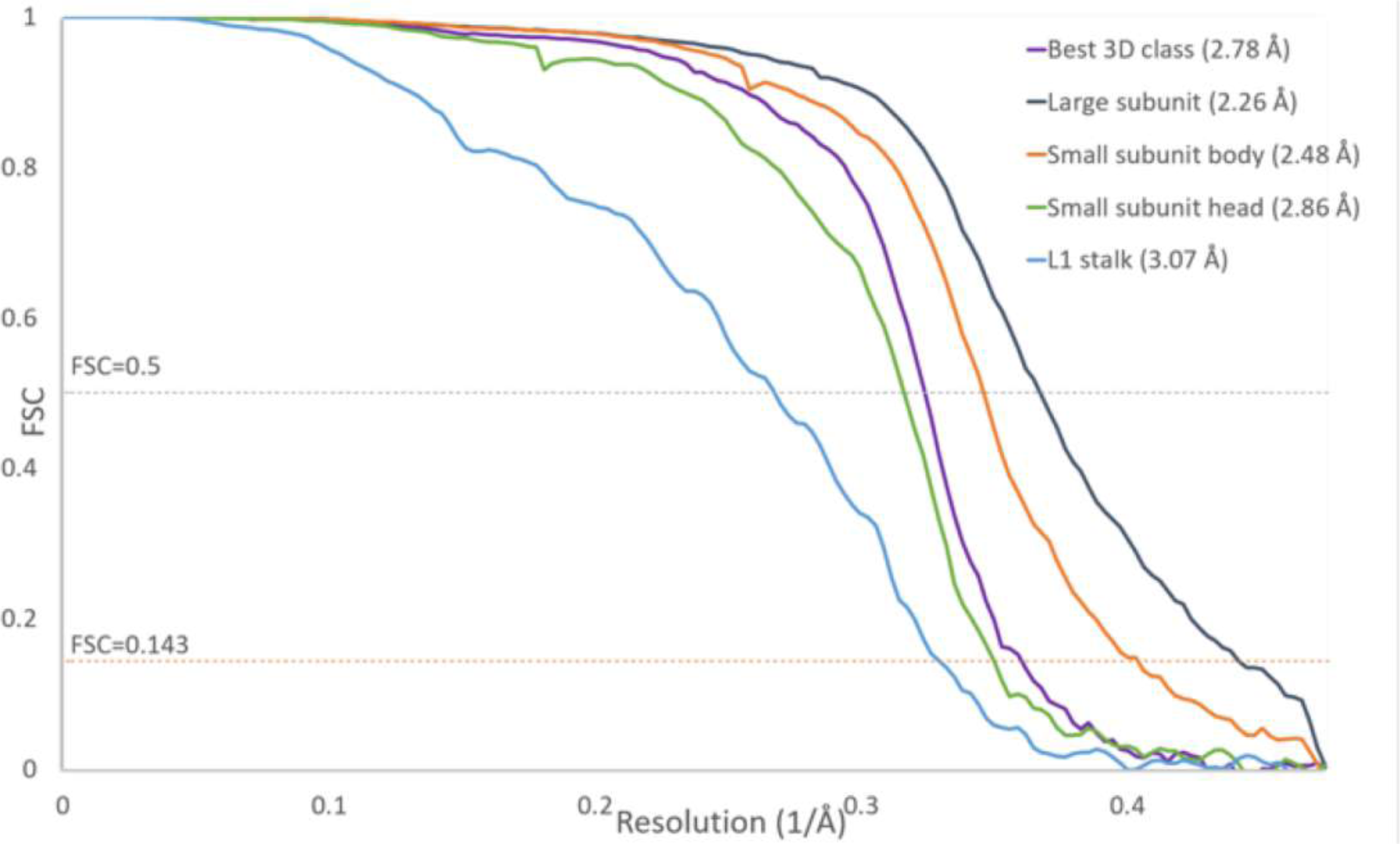
Gold standard Fourier Shell Correlation of the single particle data. Purple, best 3D class; dark blue, large subunit; orange, SSU body; green, SSU head; blue, L1 stalk.

**Figure S11.**
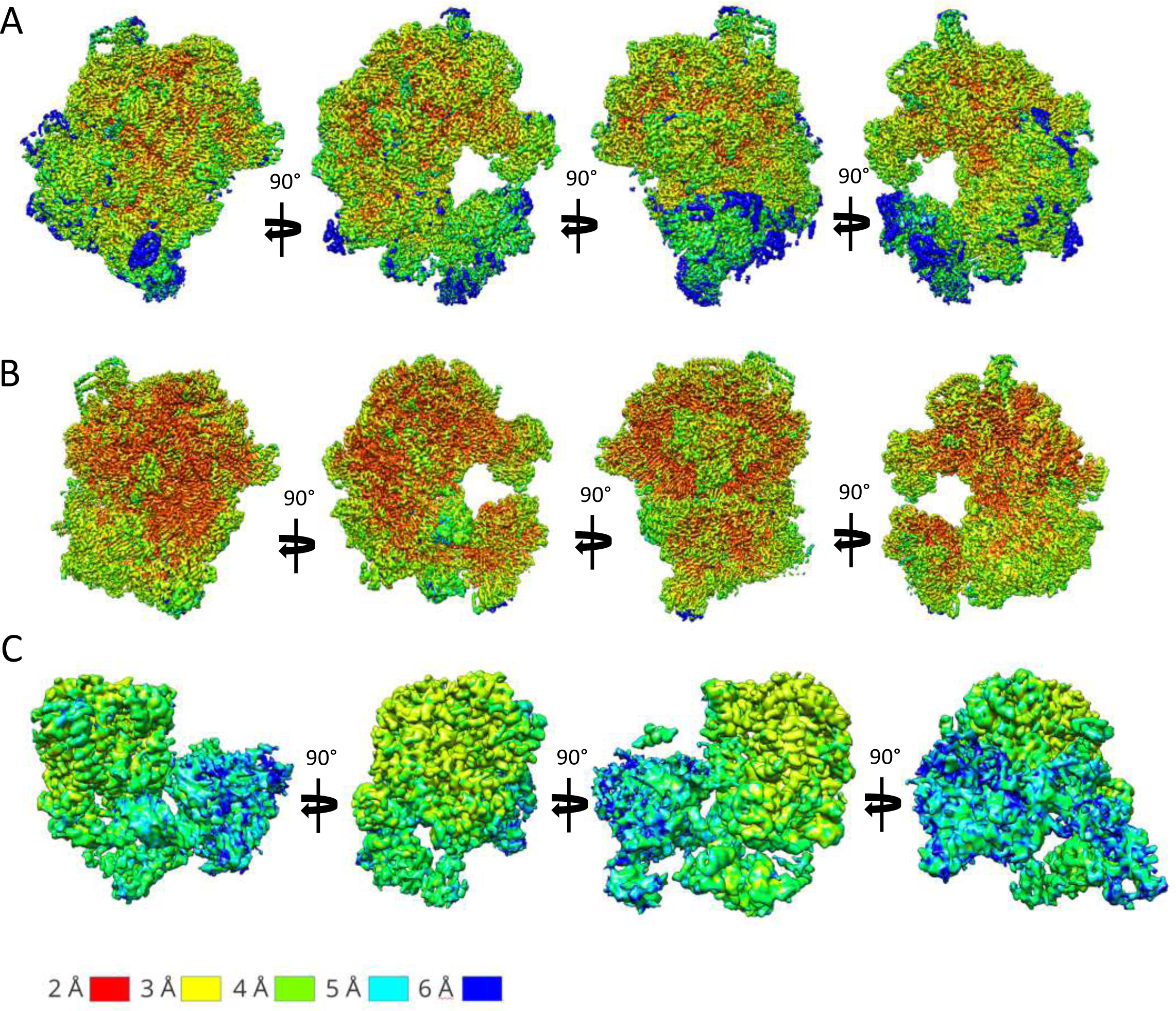
Local resolution maps of single particle analysis. **A** Whole-map refinement of the best 3D class, **B** 3-body refinement and **C** masked classification of L1 stalk.

**Figure S12.**
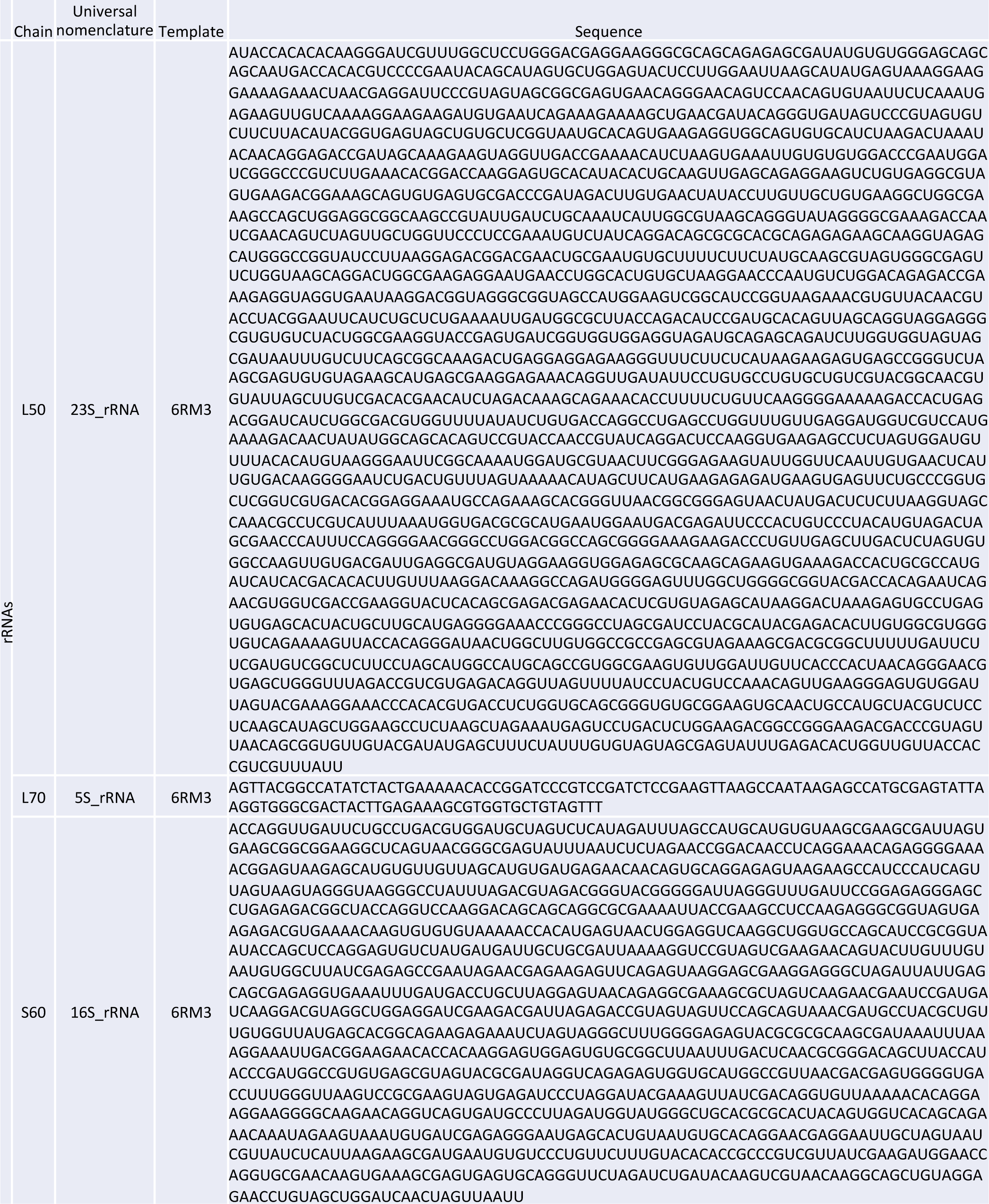
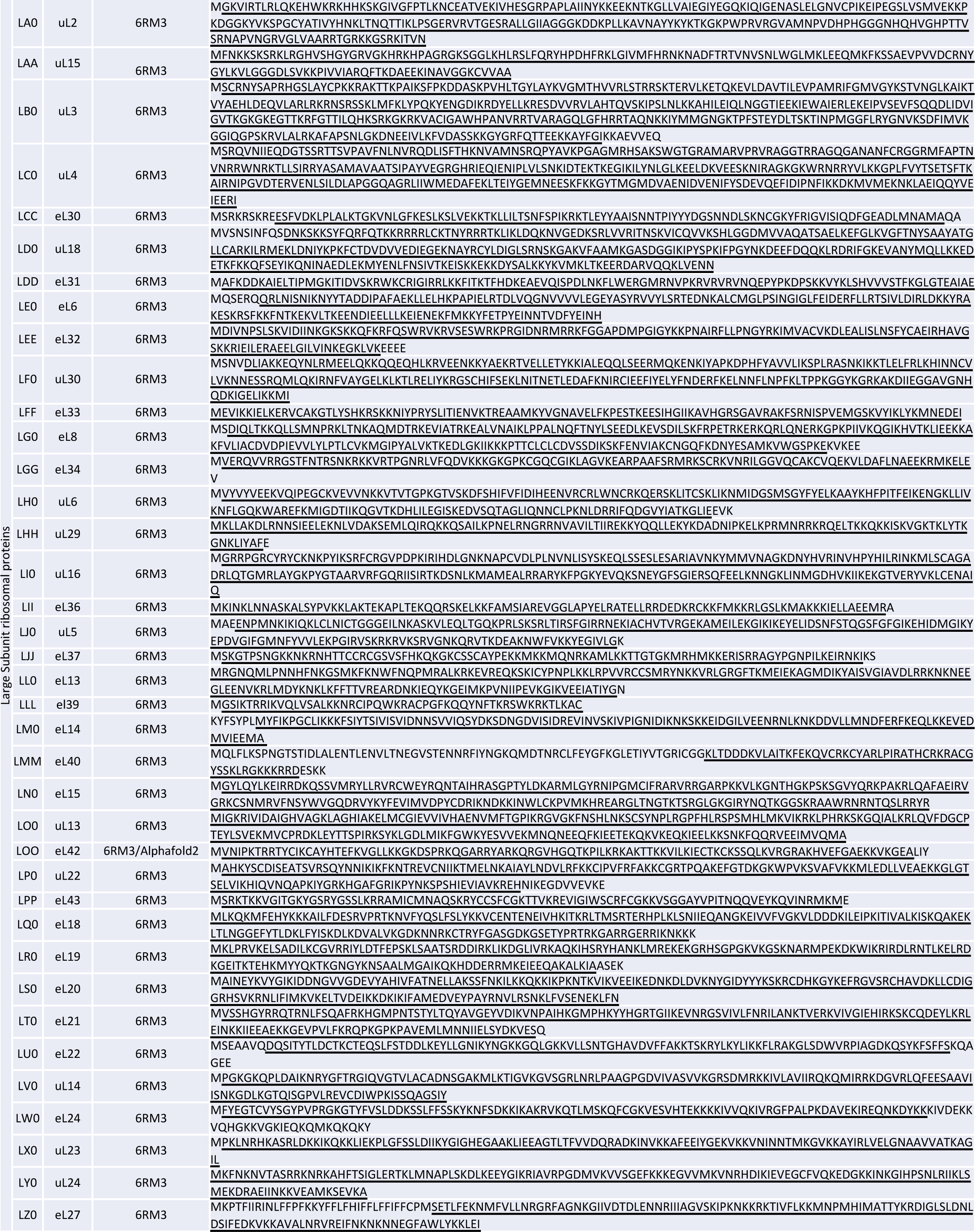
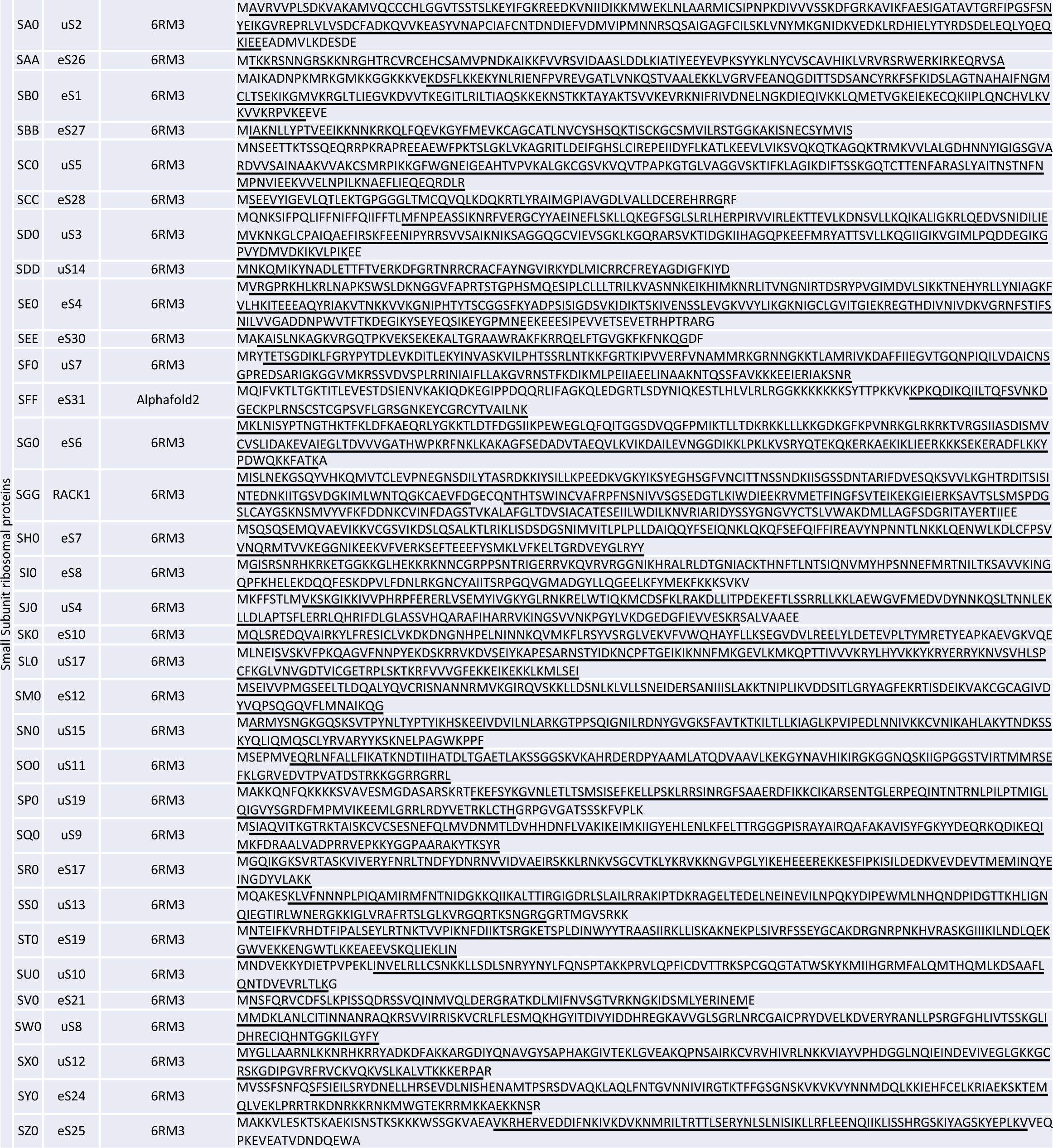
Molecular model of the *S. lophii* monomeric ribosome. Individual rRNA and protein chain sequences are listed, along with their ribosomal universal nomenclature and the templates used for model building. Protein sequences modelled in the map are underlined.

**Figure S13.**
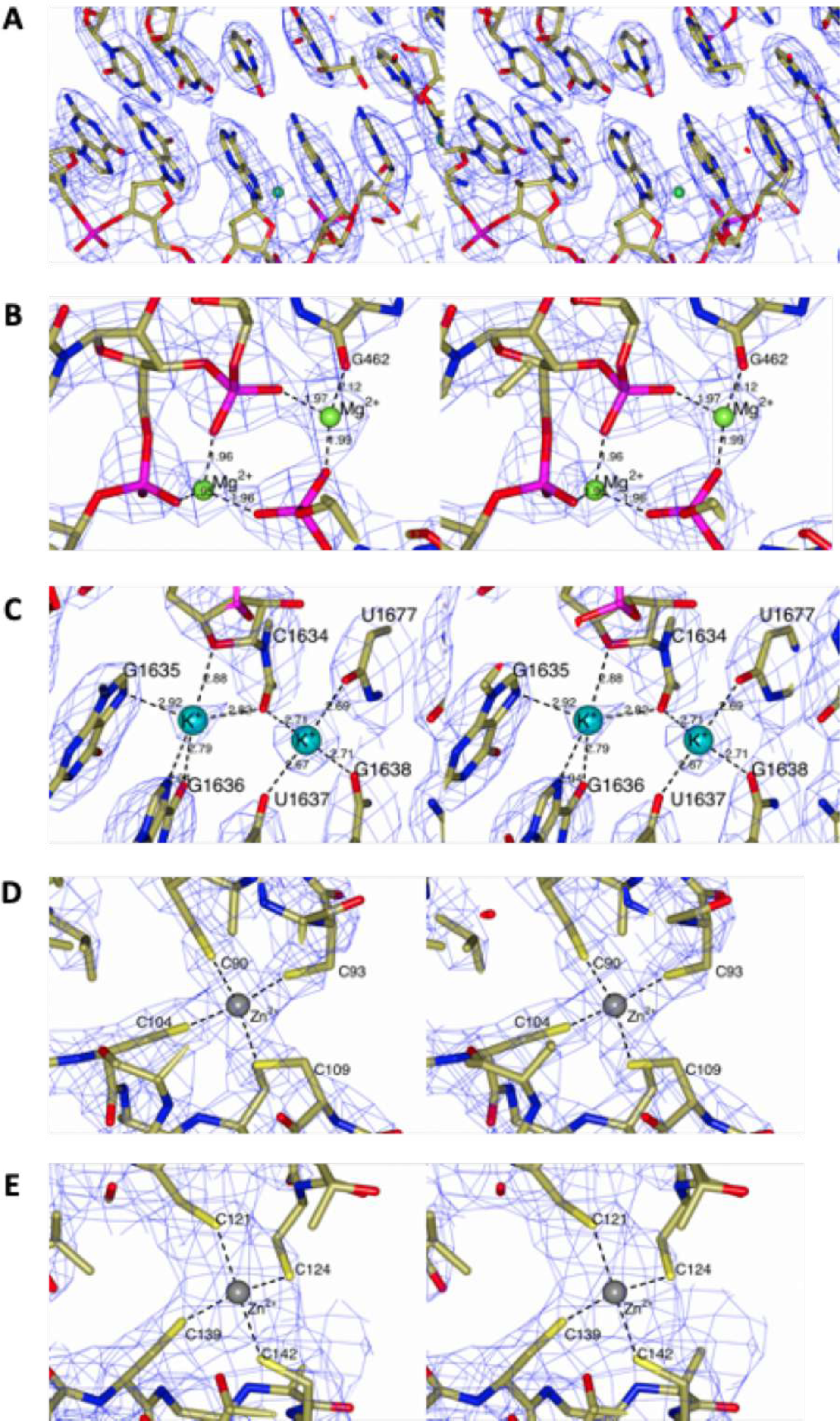
Representative closeups of the cryoEM density. Stereo diagrams showing sections of the cryo-EM density for regions of the whole-ribosome map (A-D) and SSU head map (E). **A** A representative section in the vicinity of G146 of L50 rRNA clearly shows the difference between adenine and guanine bases. **B** Density peaks are modelled as Mg2^+^ ions on the basis of short (2.0 - 2.1 Å) coordinating distances to oxygen ligands. **C** K^+^ ions are assigned on the basis of longer coordinating distances (2.6 - 2.9 Å) which may include nitrogen ligands. **D-E** Zn^2+^ ion modelling in eL40 and eS31 chains.

**Figure S14.**
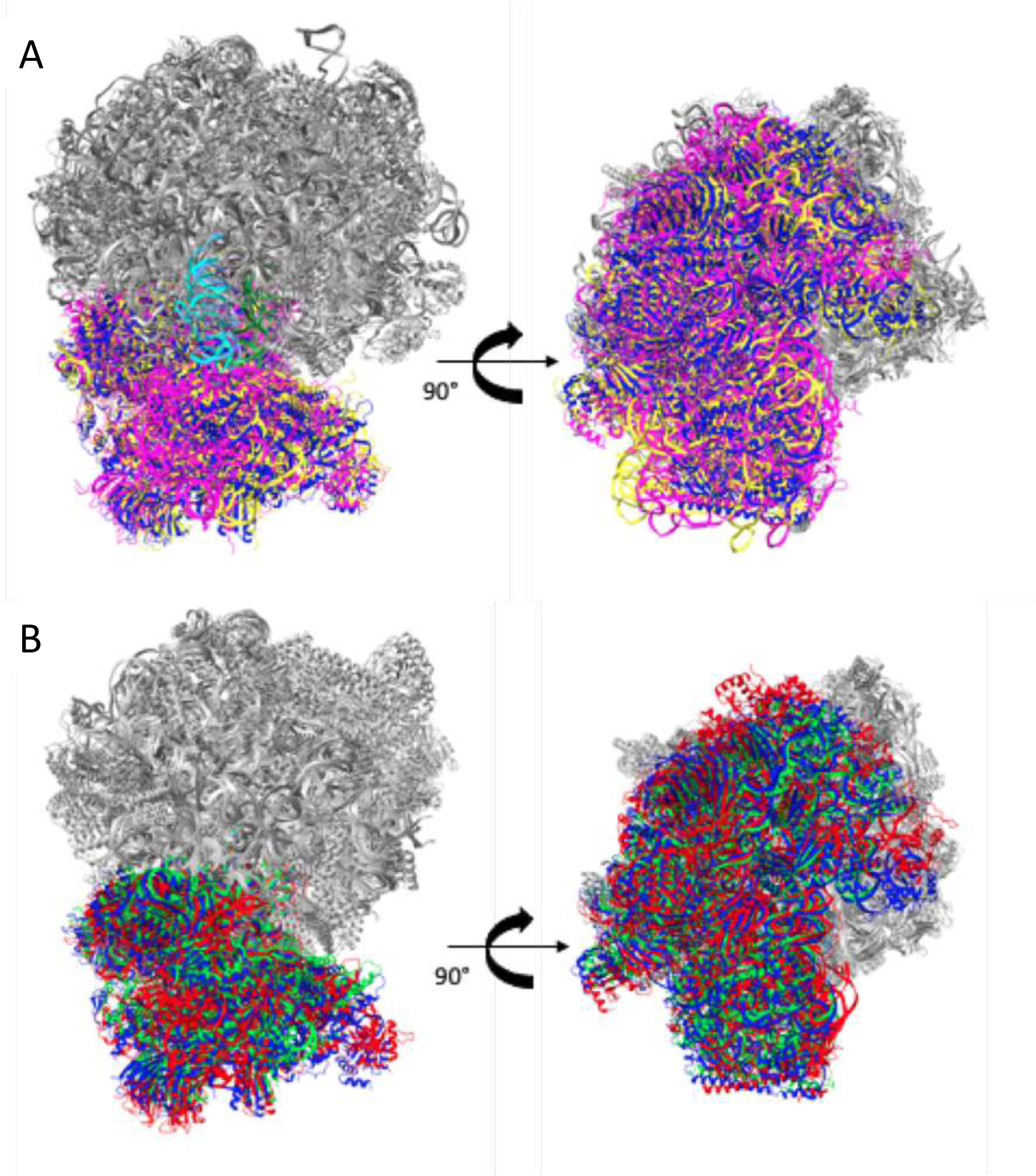
The *S. lophii* ribosome is in a non-rotated state. **A** *S. lophii* ribosome structure (7QCA) shown in ribbon representation superimposed with *S. cerevisiae* ribosome structures in rotated (3J77) and non-rotated (3J78) states. Large subunits are coloured grey and small subunits blue (*S. lophii*), magenta (*S. cerevisiae*, rotated state) and yellow (*S. cerevisiae*, non-rotated state). tRNA molecules bound to the yeast non-rotated state are coloured in cyan and green. Ribosomes are shown in a side view in the left-hand panel, and then rotated by 90° to show a bottom view in the right-hand panel. **B** *S. lophii* ribosome structure (7QCA) shown in ribbon representation overlaid with other microsporidian ribosome structures. Large subunits are coloured grey and small subunits blue (*S. lophii*), red (*V. necatrix*, rotated state) and green (*P. locustae*, non-rotated state). Ribosomes are shown in a side view in the left-hand panel, and then rotated by 90° to show a bottom view in the right-hand panel.

**Figure S15.**
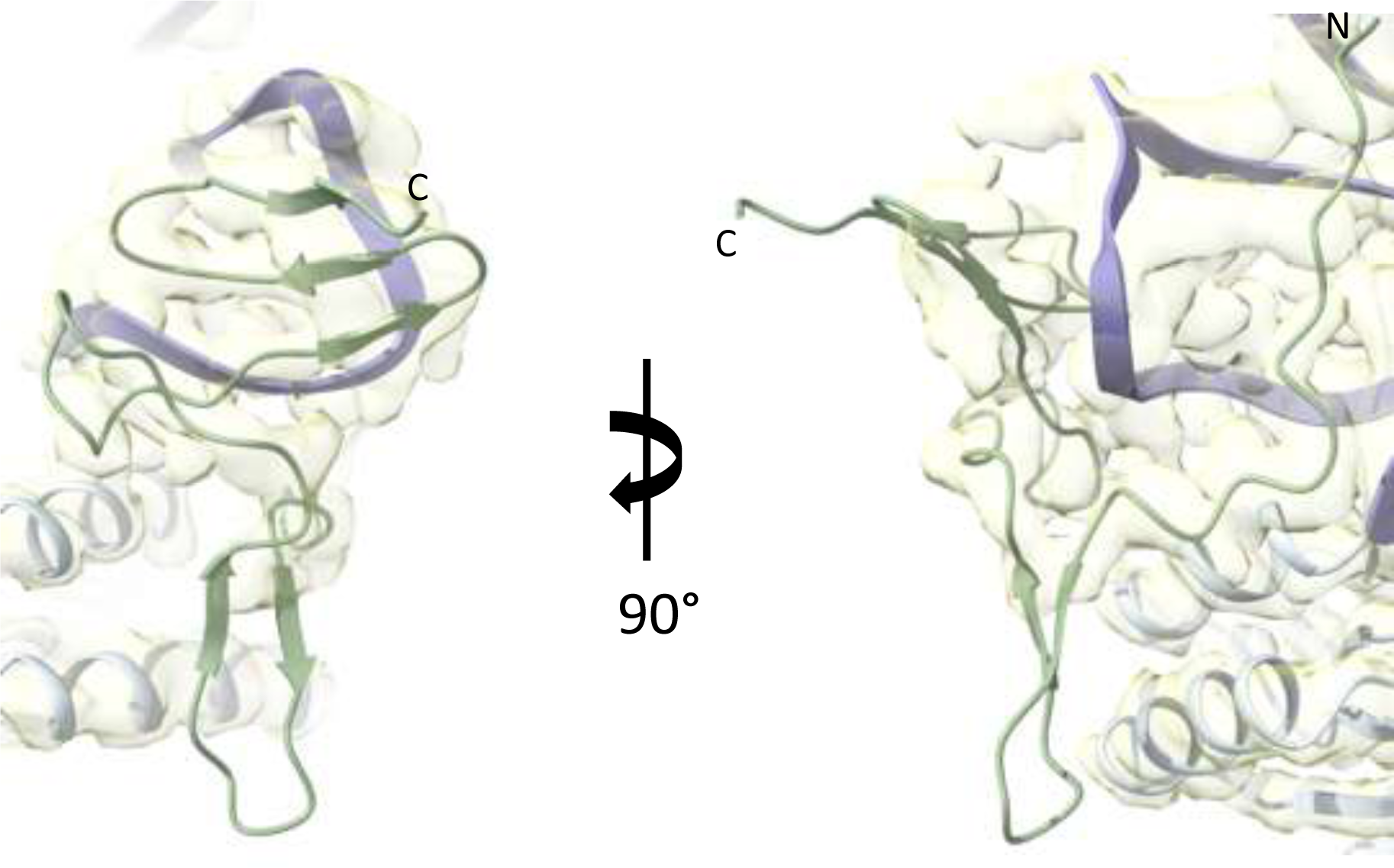
Atomic model of subunit eS31. The structure of *S. lophii* eS31 (green ribbon) was predicted in Alphafold2 and refined against the cryoEM map of the *S. lophii* ribosome (transparent yellow). The neighbouring 16 S rRNA (purple ribbon) and the protein subunit eS12 are also shown.

**Figure S16.**
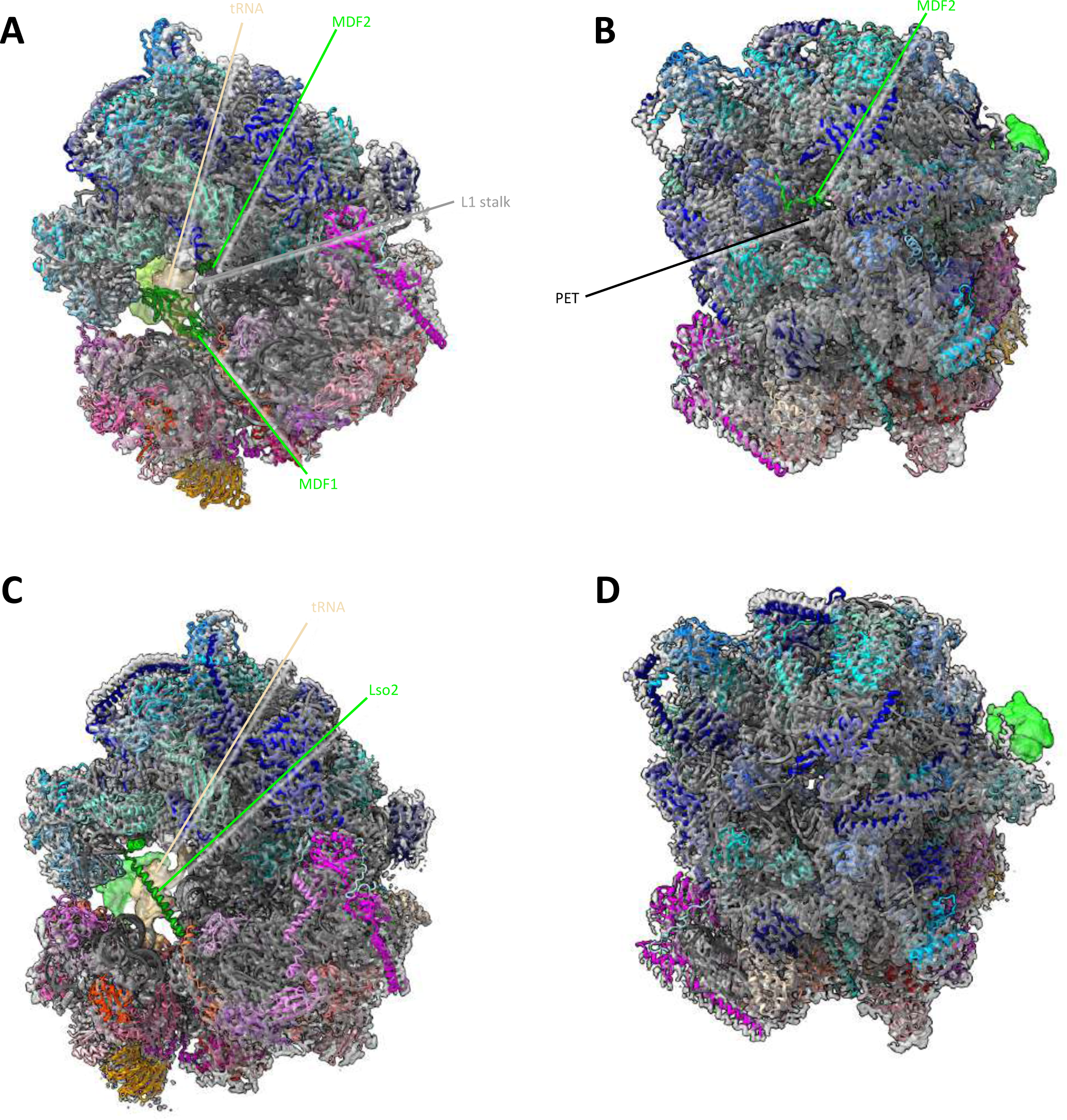
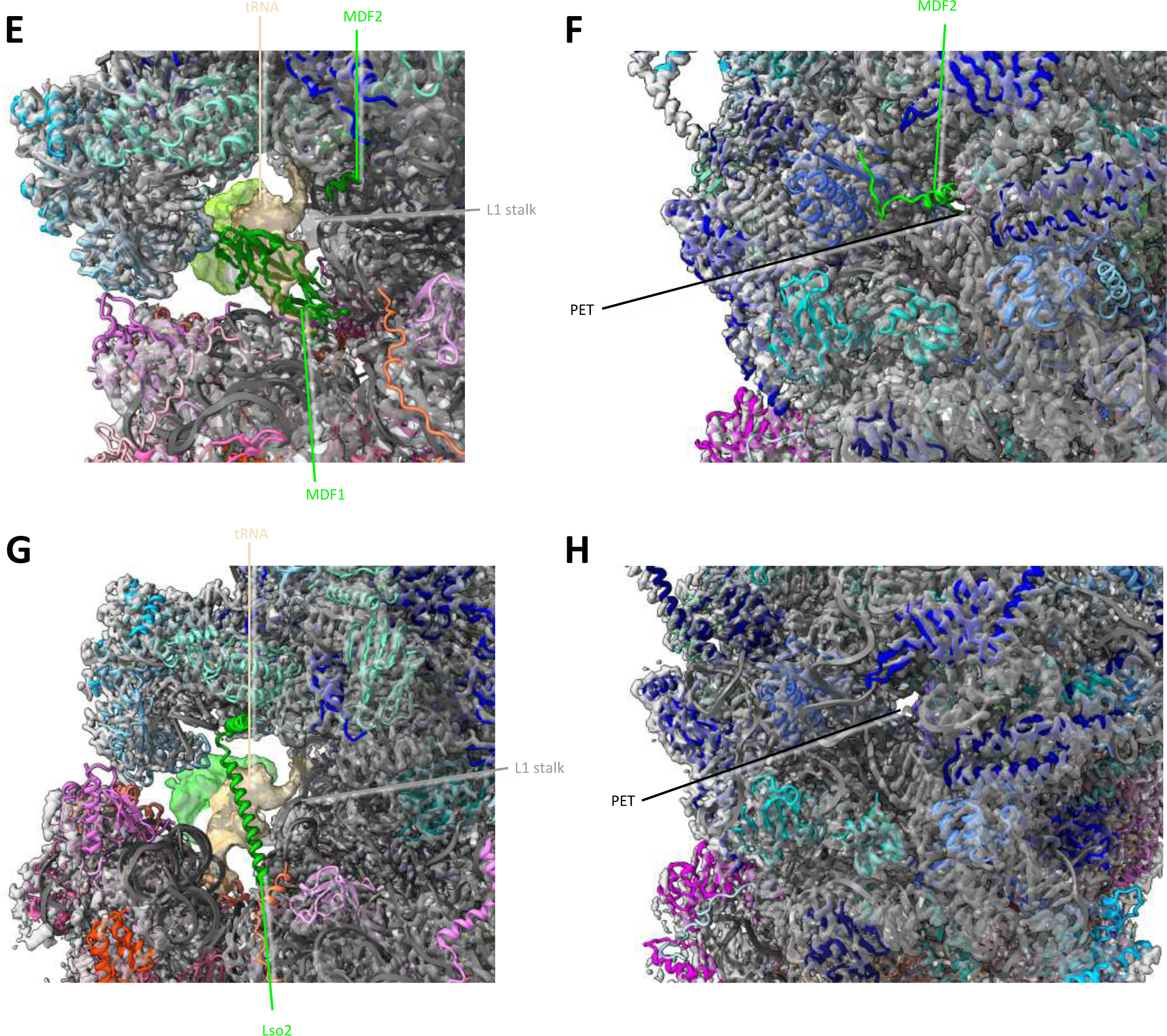
Absence of hibernation factors in the *S. lophii* single particle map. The single particle map of the *S. lophii* ribosome shows no densities for hibernation factors found in the microsporidian species *V. necatrix* and *P. locustae.* **A, B** Atomic model of the *V. necatrix* ribosome (multicolour) fitted into the density map of *S. lophii* (transparent grey). The hibernation factor MDF1 blocking the mRNA and tRNA binding sites (green ribbon) is shown in A and the hibernation factor MDF2 occupying the polypeptide exit tunnel (PET) is shown with the same colour in B. **C, D** Atomic model of the *P. locustae* ribosome (multicolour) fitted into the density map of *S. lophii* (transparent grey). The hibernation factor Lso2 is shown as a green ribbon, occupying the tRNA binding sites in C and sequestering the PET in D. **E-H** Closeups of A-D.

**Figure S17.**
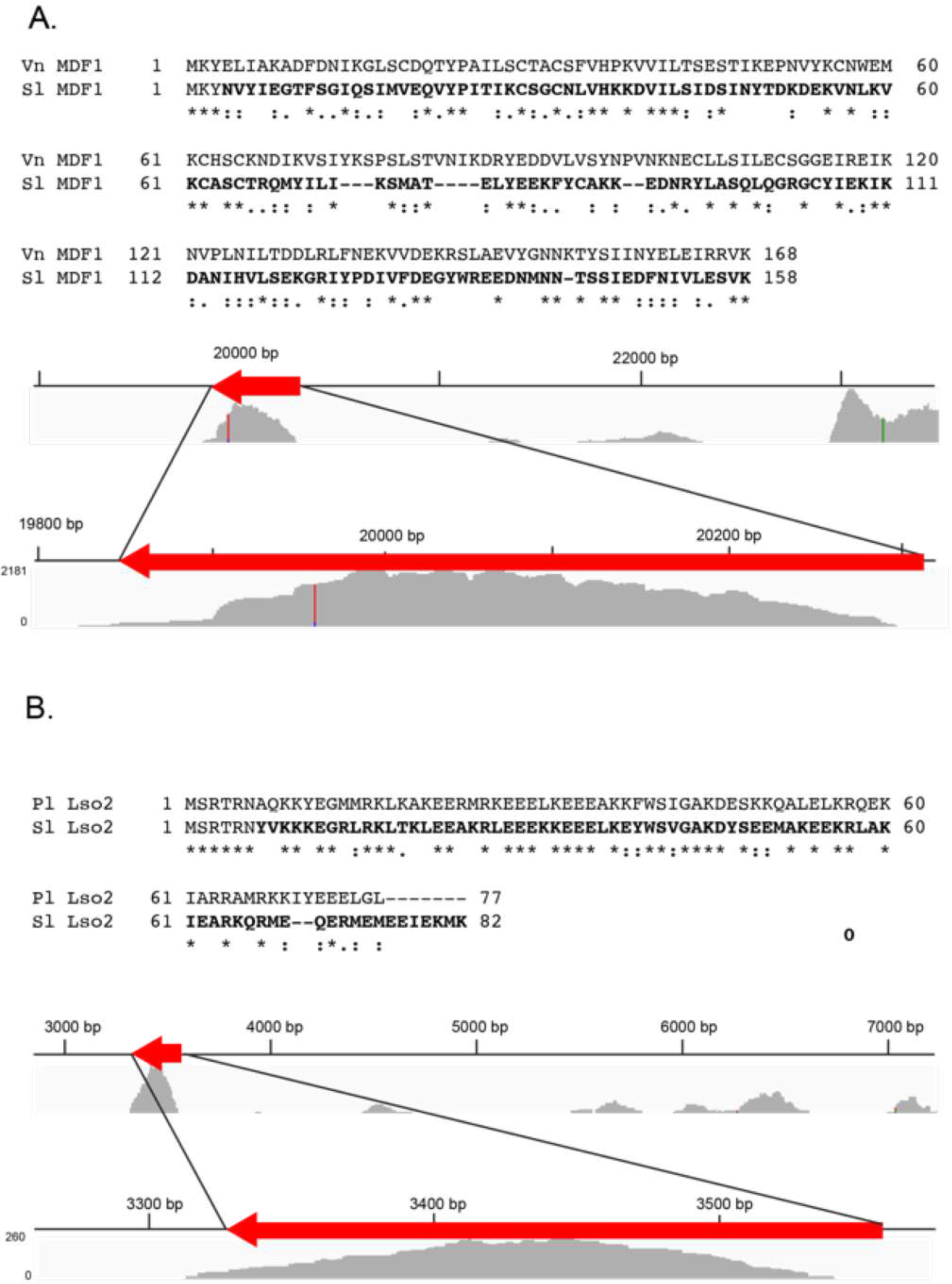
MDF1 and Lso2 homologs are encoded in the genome of *S. lophii*. Sequence alignments of ribosomal hibernation factors. **A** BLASTP search of the *S. lophii* protein database with the MDF1 protein sequence from *V. necatrix* resulted in a hit for hypothetical protein SLOPH_1662 (EPR78075.1) with 29% sequence identity and 51% sequence similarity. **B** A putative Lso2 protein was identified in the *S. lophii* genome on contig ATCN01000495.1 using TBLASTN with the Lso2 from *P. locustae*. The sequences have 52% identity and 68% similarity. RNASeq reads from the *Spraguea lophii* transcriptome (SRX312311) were mapped onto the Celtic Deep *Spraguea lophii* assembly (GCA_001887945.1) using Bowtie2^53^ with default settings. Upper contigs show the context of the genes of interest and RNAseq coverage compared to immediately adjacent regions of the genome and lower contig regions focus on RNAseq coverage of the gene of interest.

**Figure S18.**
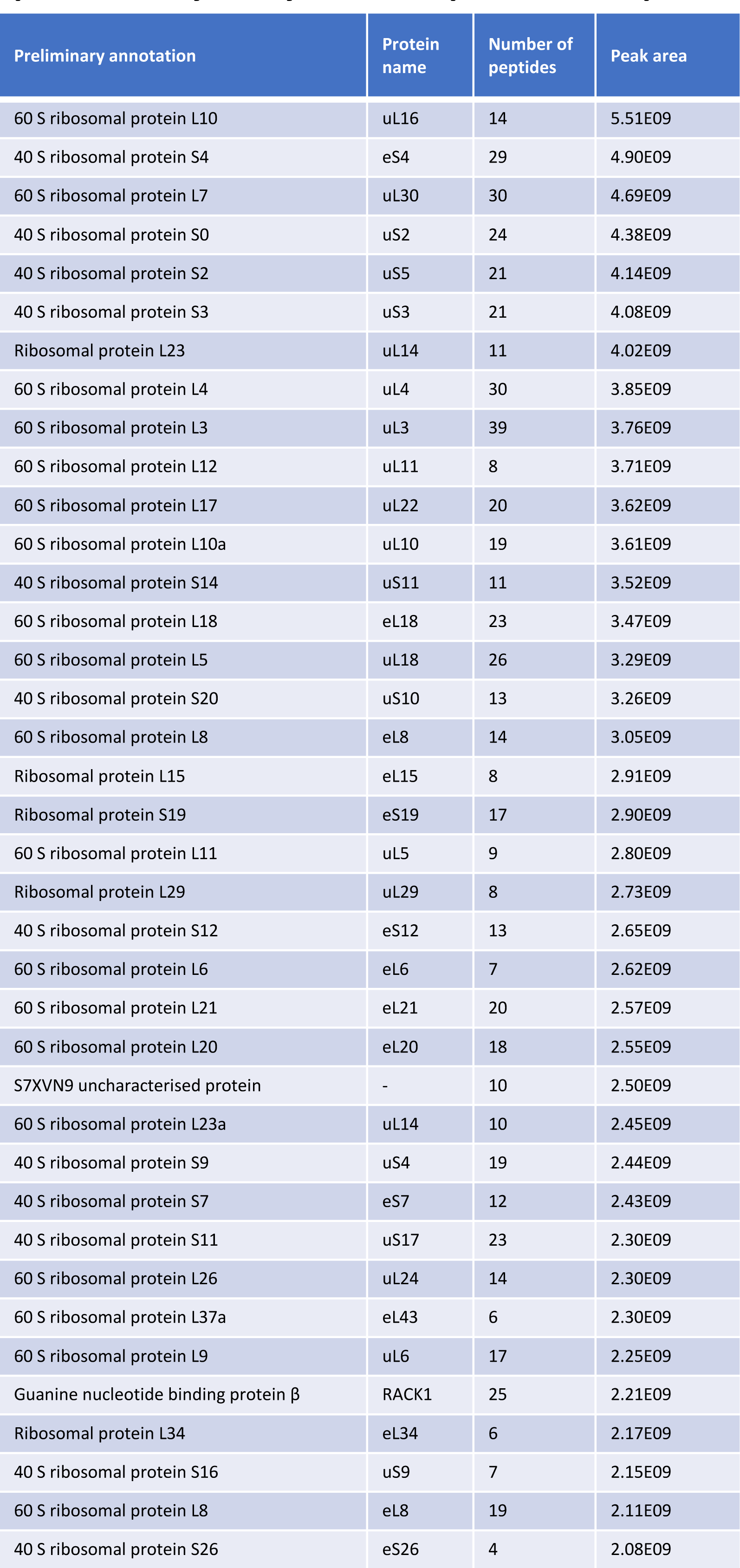
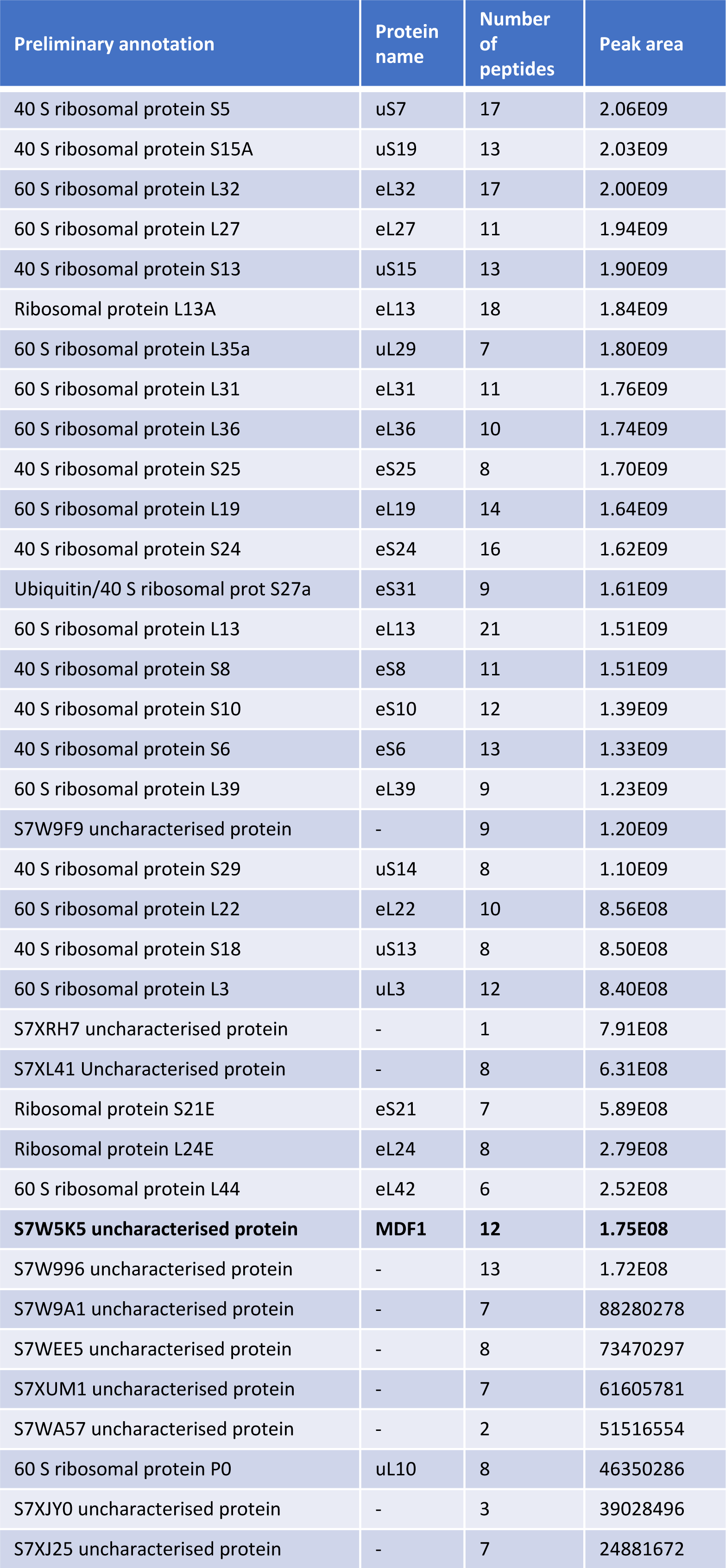
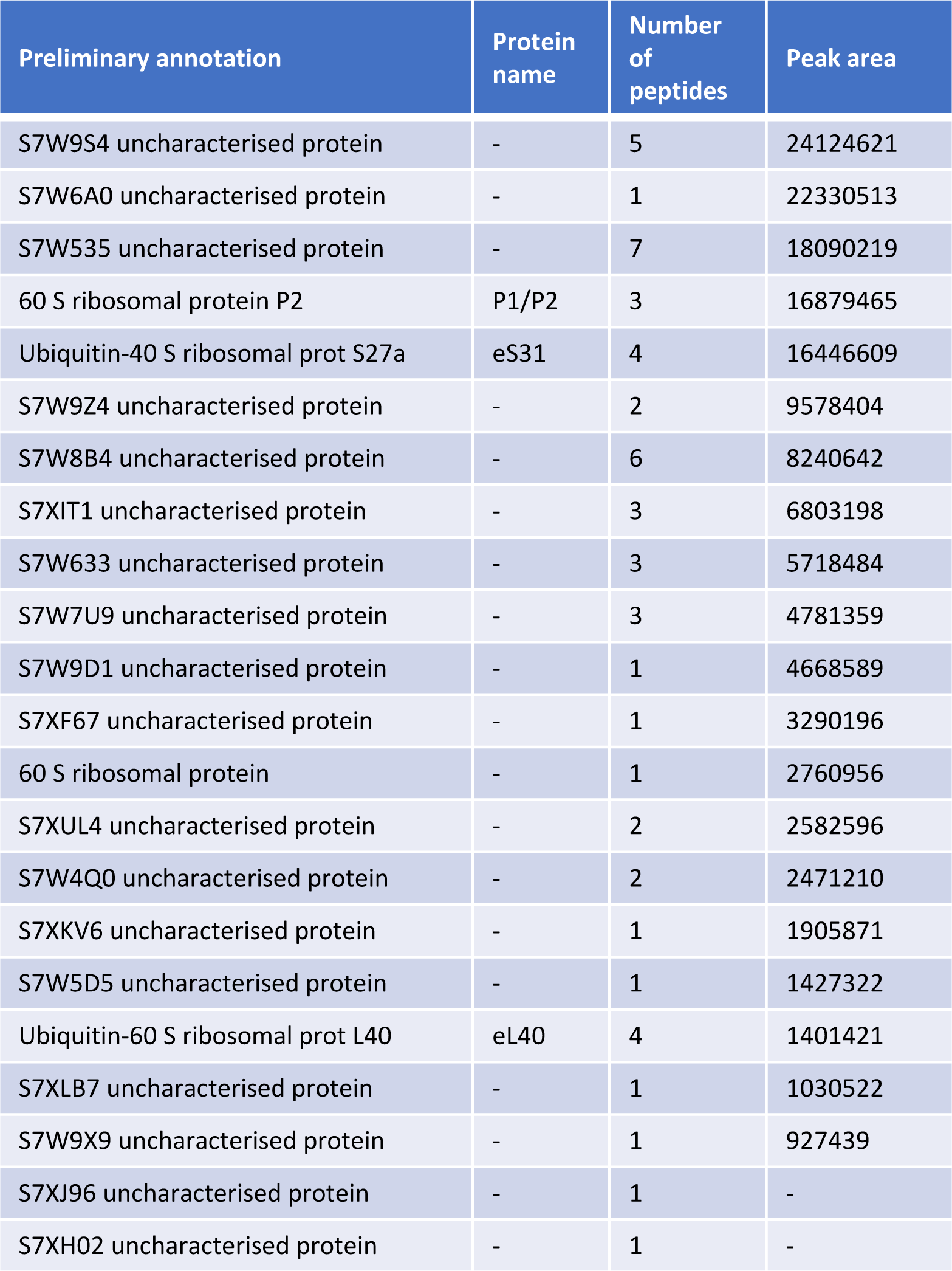
Mass spectrometry analysis of the purified *S. lophii* ribosome sample. List of ribosomal and uncharacterised proteins identified in the mass spectrometry analysis. The microsporidian hibernation factor MDF1 is highlighted in bold. As an approximation of protein abundances in the sample, the peak area was used^16^ and the proteins are presented in peak height order.

**Figure S19.**
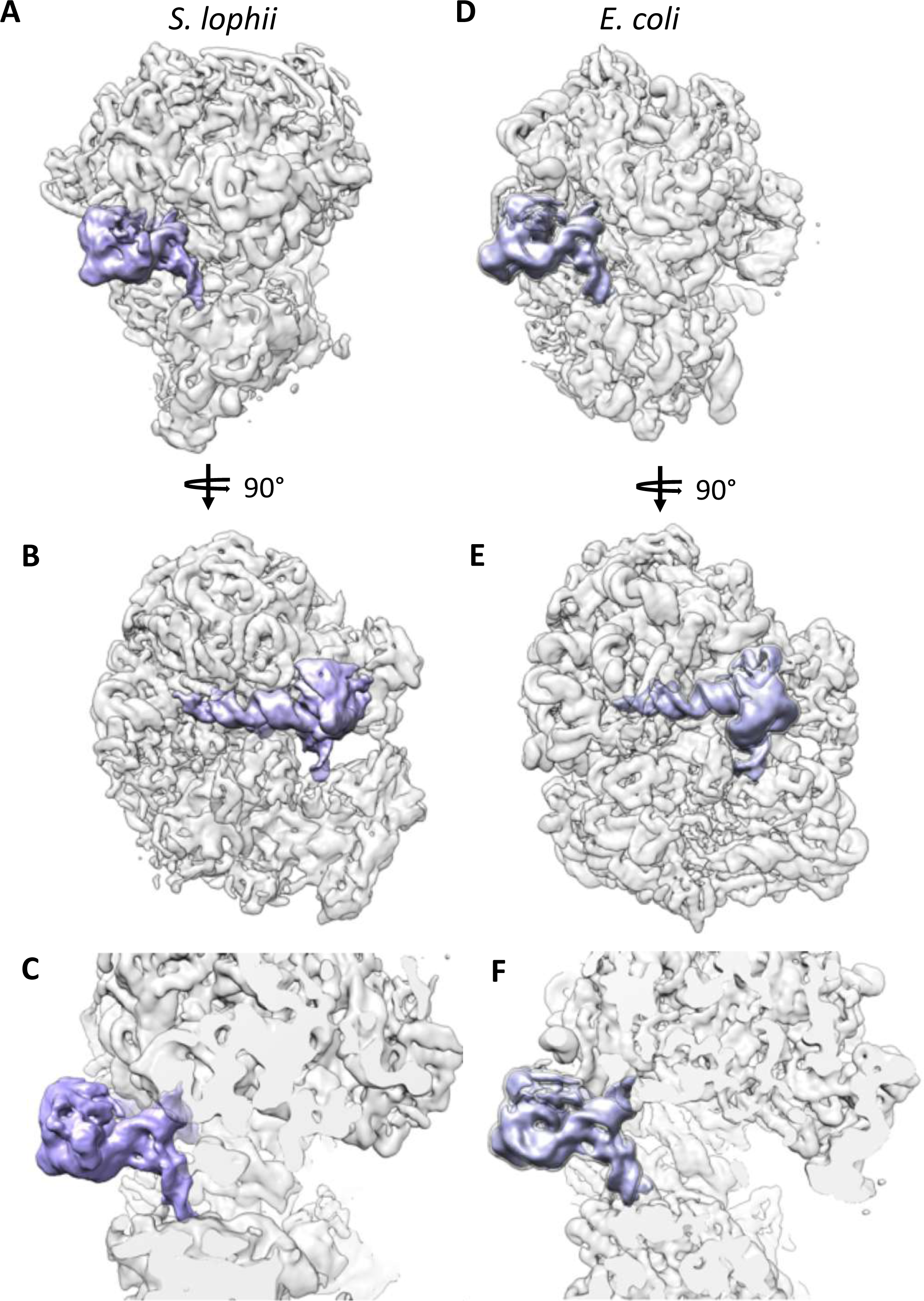
The L1 stalk and E site tRNA in the hibernating *E. coli* and *S. lophii* ribosomes. The hibernating ribosome from *E. coli* (EMD-0137)^9^ (**A-C**) compared with the hibernating *S. lophii* ribosome (**D-F**). Both cryoEM maps (transparent) were filtered with a positive B-factor of 200 Å^2^ for clarity and the density of the L1 stalk and the bound tRNA are highlighted in purple. **B** and **E** are rotated by 90 degrees with respect to **A** and **D,** and **C** and **F** are clipped closeups of A and D, respectively. The structure and orientation of the L1 stalk and bound E site tRNA are virtually the same in both structures.

**Figure S20.**
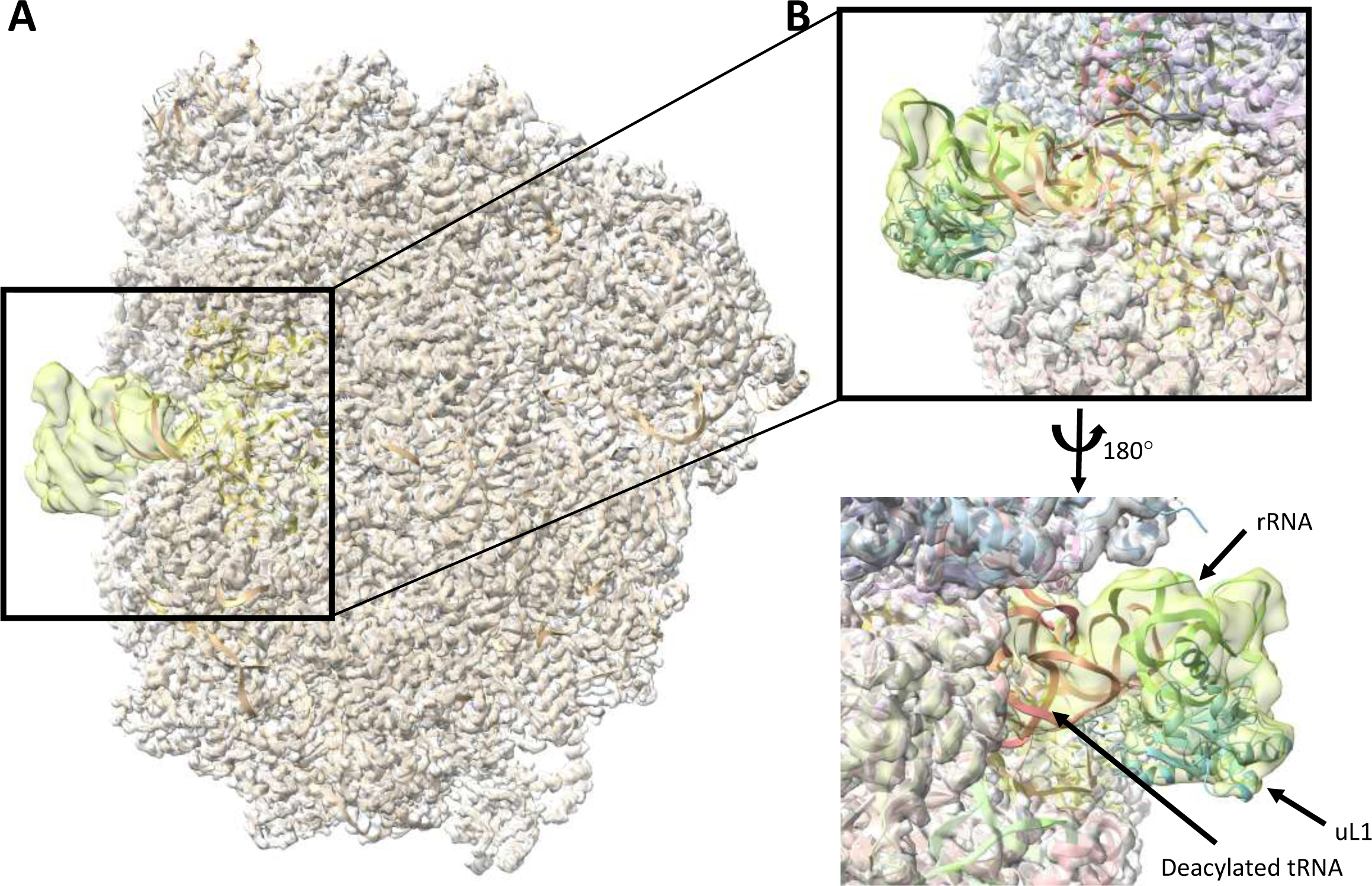
Atomic model of the L1 stalk. **A** The density map of the whole ribosome is shown as a grey surface with the model as peach ribbons. The map of the L1 stalk (filtered with a B factor of 200 Å^2^) is shown as a yellow surface. **B** A close up of the L1 stalk showing the modelled rRNA as a green ribbon, protein uL1 as a cyan ribbon and a deacylated tRNA in the ribosome E site as an orange ribbon.

**Figure S21.**
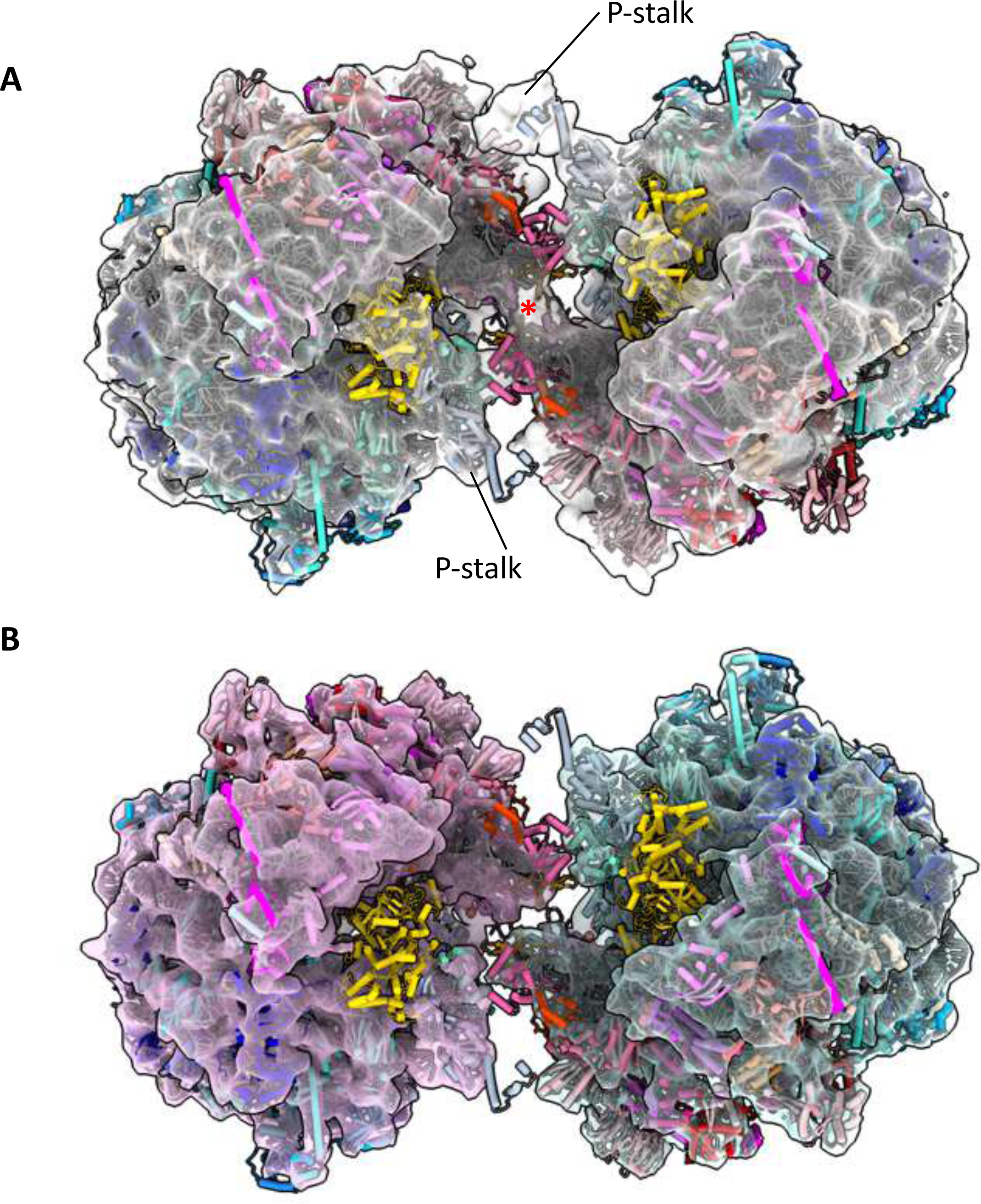
Atomic model of the *S. lophii* 100 S ribosome. **A** Two atomic models of the 70 S ribosome fitted into the *in situ* 15.2 Å sub-tomogram average of the 100 S dimer. Red asterisk, bridging density between the subunits eS31 and rRNA of the SSU. **B** Two copies of the 10.3 Å sub-tomogram average obtained by calculating for one half of the dimer arranged as in A.

**Figure S22.**
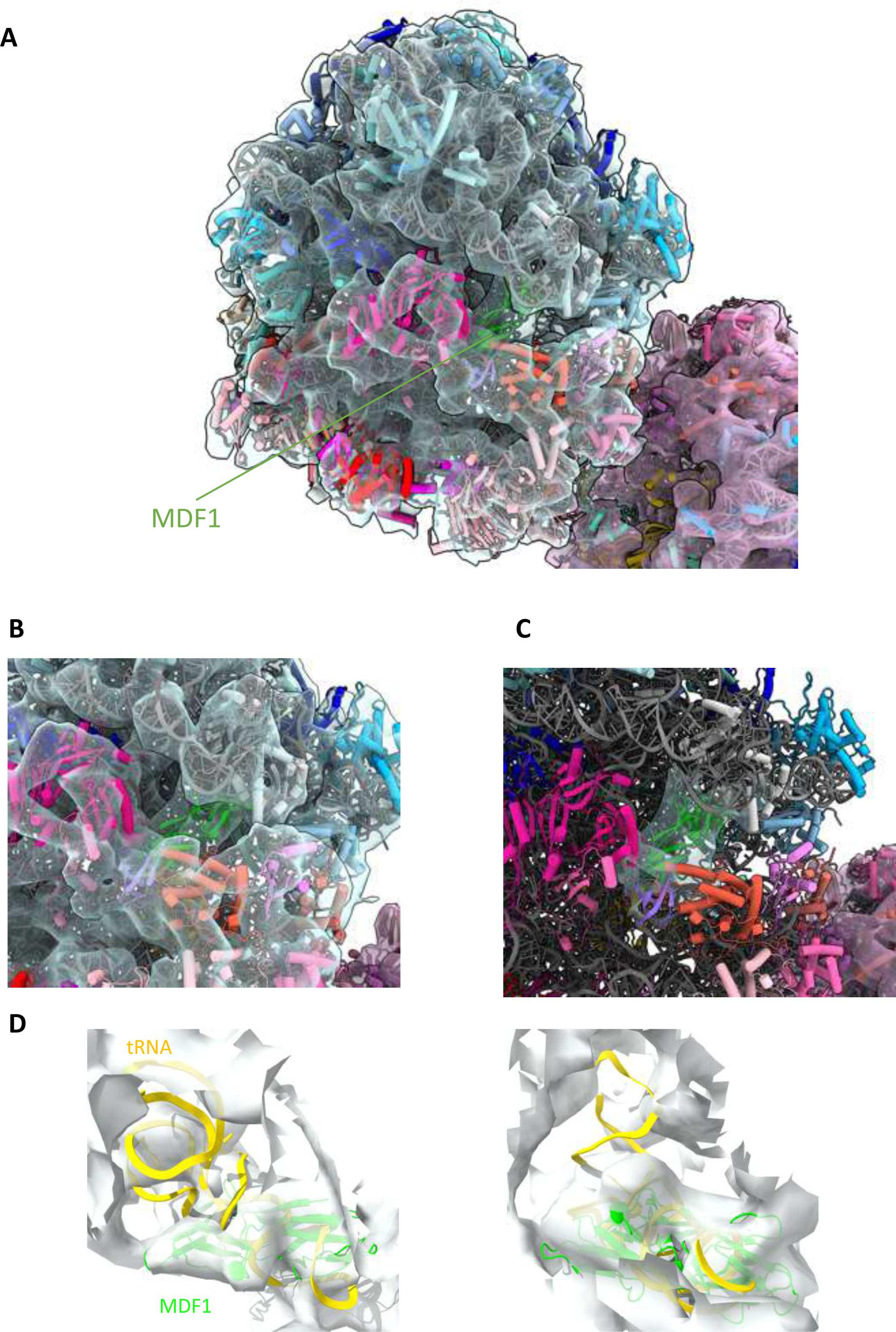
Evidence for MDF1 bound in the *in situ* sub-tomogram average. **A** Atomic model of the *S. lophii* ribosome in cartoon representation (colour scheme as in Fig. 2) fitted into the 10.3 Å sub-tomogram average map of the *S. lophii* dimer. Density for MDF1 (green ribbon) was observed. **B** Close up of the map showing the MDF1 binding site and **C** map is drawn as a 12 Å zone around the MDF1 molecule for clarity. **D** Two views showing the MDF1-bound dimer structure overlaid with the tRNA-bound monomer structure to highlight that there was no observed density in the dimer map for the acceptor stem of the tRNA molecule, and that MDF1 was the best fit to the map. tRNA shown in yellow, MDF1 in green, map has been drawn as an 18 Å zone around the tRNA molecule for clarity.

**Figure S23.**
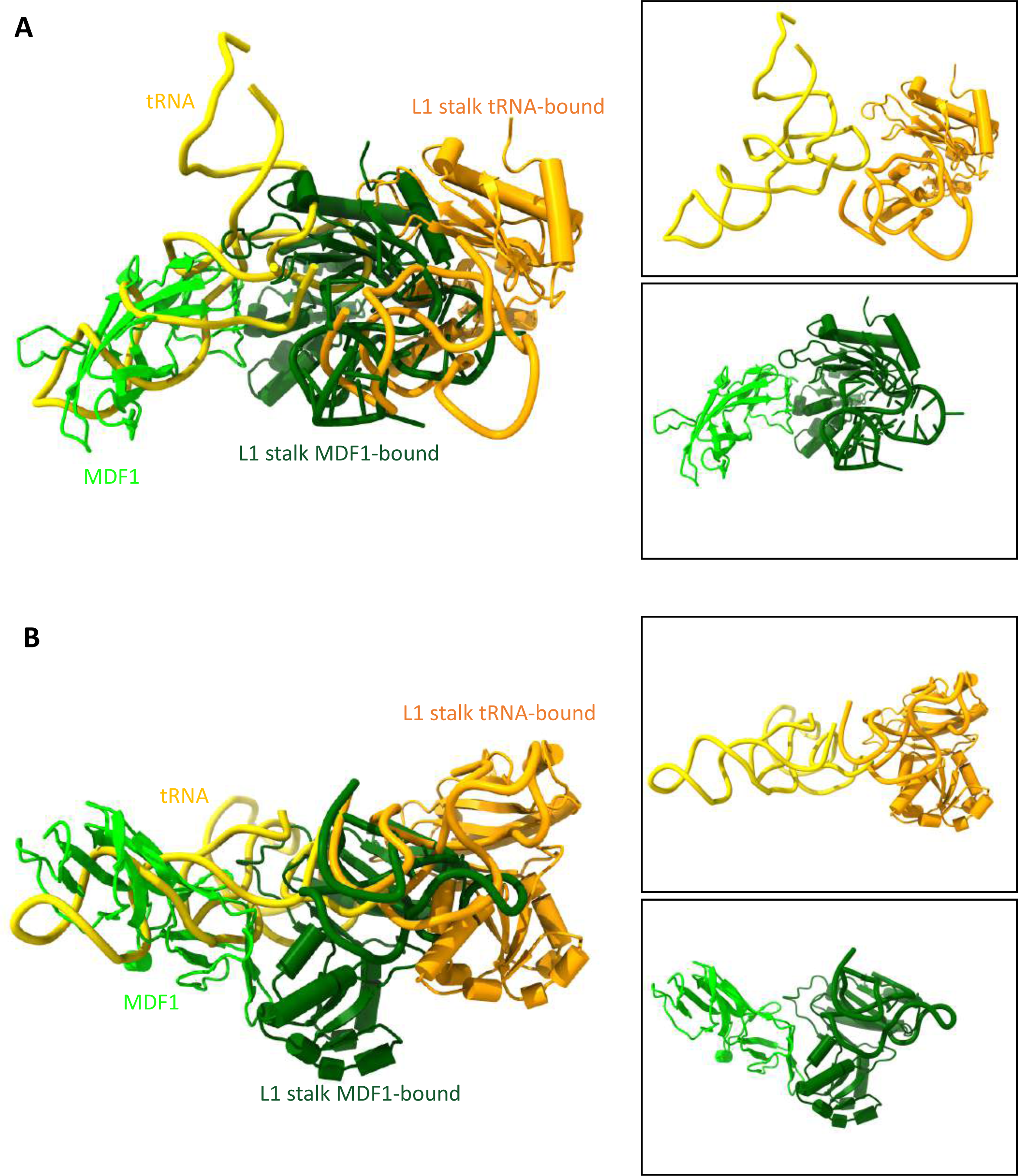
Comparison of the L1 stalk position in the tRNA-bound and MDF1-bound states. **A and B** Two different views of the L1 stalk in an overlay of the tRNA-bound and MDF1-bound states. tRNA shown in yellow with corresponding L1 stalk (consisting of the uL1 protein and rRNA) in orange (from the monomeric ribosome structure). MDF1 in lime with corresponding L1 stalk (consisting of the uL1 protein and rRNA) in dark green (from the dimeric ribosome structure). Boxed figures show the tRNA-bound and MDF1-bound states alone for each view.

**Figure S24.**
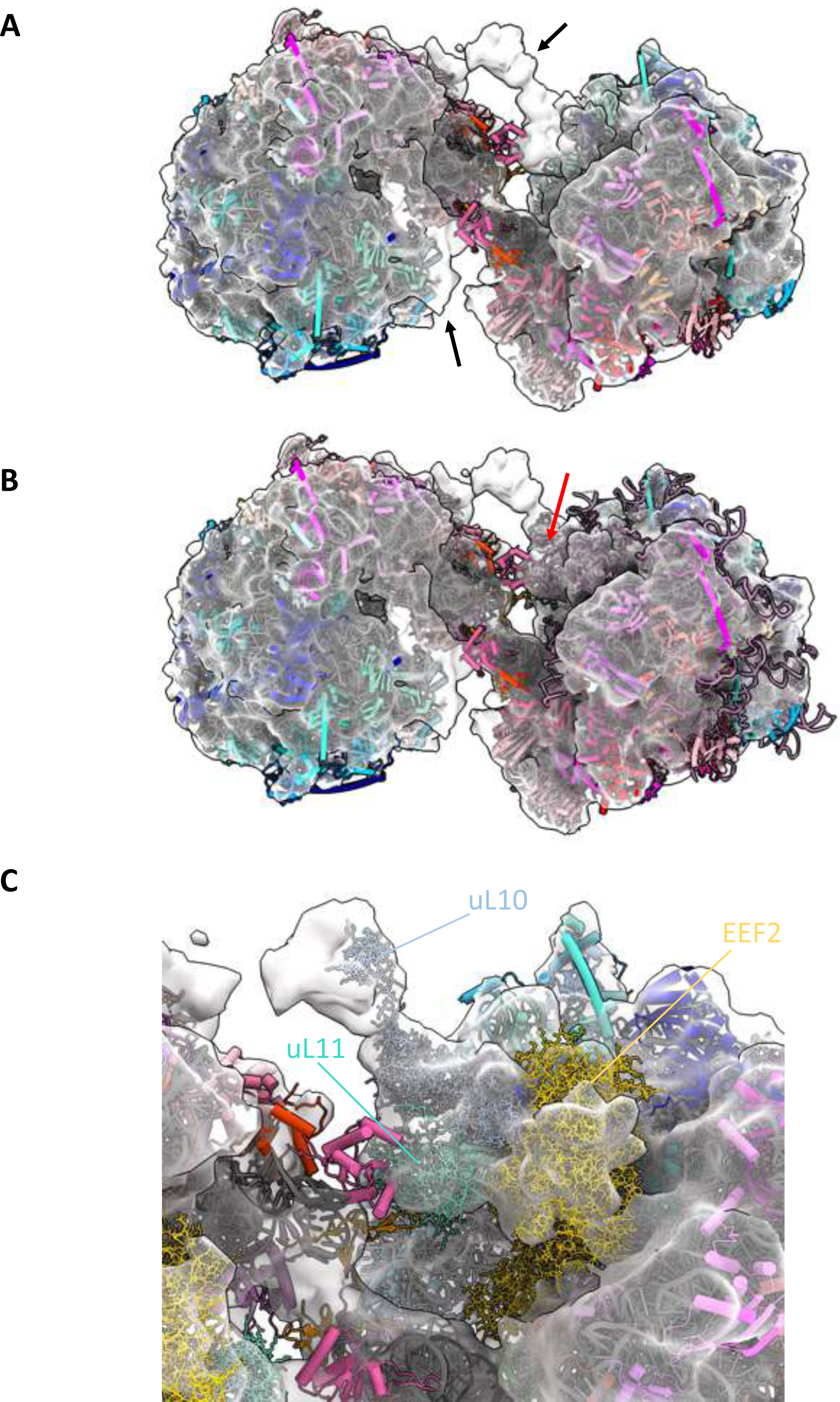
The P-stalk density of the 100 S *S. lophii* ribosome. **A** *S. lophii* dimer model (cylinders and stubs) and map (transparent surface) showing density for the P-stalk (black arrow). **B** Structure of the ribosome from pig (3J7P^44^; purple ribbons) overlaid on one of the *S. lophii* ribosomes. Red arrow shows some of the P-stalk density is filled. **C** Alphafold2 models of the S. lophii proteins uL10 (blue sticks), uL11 (cyan sticks) and eukaryotic elongation factor 2 (gold sticks) were modelled into the dimer map guided by the structure of the pig ribosome.

**Figure S25.**
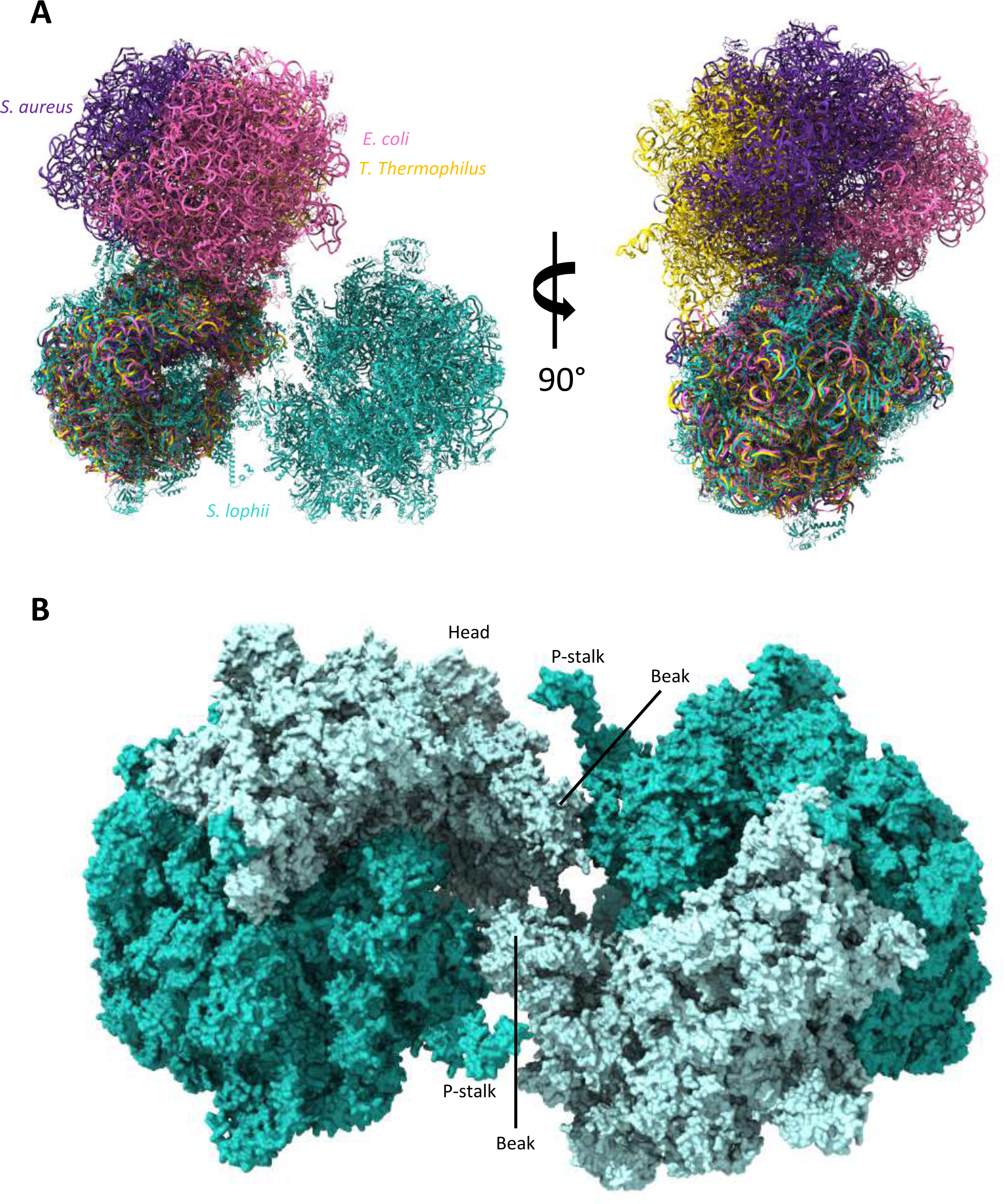
Comparison of *S. lophii* dimer and bacterial dimers. **A** Overlay of *S. lophii* ribosome dimer (light sea green) with bacterial ribosomes dimers from *S. aureus* (6FXC, purple), *T. themophilus* (6GZX, gold) and *E. coli* (6H58, pink) shown in two views, rotated by 90°. Structures were overlaid using one subunit of the dimer. **B** The *S. lophii* dimer structure in surface representation with labelled parts showing the dimerization interface is located between the beak regions of the small subunits.

